# Stoichiometric Analysis of Protein Complexes by Cell Fusion and Single Molecule Imaging

**DOI:** 10.1101/2020.01.26.920058

**Authors:** Avtar Singh, Alexander L. Van Slyke, Maria Sirenko, Alexander Song, Paul J. Kammermeier, Warren R. Zipfel

**Affiliations:** Applied and Engineering Physics, Cornell University, Ithaca, NY 14853; Graduate Field of Biophysics, Cornell University, Ithaca, NY 14853; Department of Biological Engineering, Cornell University, Ithaca, NY 14853; Department of Pharmacology and Physiology, University of Rochester Medical Center, Rochester, NY; Meinig School of Biomedical Engineering, Cornell University, Ithaca, NY 14853

**Keywords:** Single Molecule Imaging, FCS, Protein complex stoichiometry, VSVG, Cell fusion

## Abstract

The composition, stoichiometry and interactions of supramolecular protein complexes are a critical determinant of biological function. Several techniques have been developed to study molecular interactions and quantify subunit stoichiometry at the single molecule level; however, these typically require artificially low expression levels to achieve the low fluorophore concentration required for single molecule imaging, or use of detergent isolation of complexes that may perturb native subunit interactions. Here we present an alternative approach where protein complexes are assembled at physiological concentrations and subsequently diluted *in situ* for single-molecule level observations while preserving them in a near-native cellular environment. We show that coupling this *in situ* dilution strategy with single molecule techniques such as *in vivo* Fluorescence Correlation Spectroscopy (FCS), bleach step counting for quantifying protein complex stoichiometry, and two-color single molecule colocalization, improves the quality of data obtained using these single molecule fluorescence methods. Single Protein Recovery After Dilution (SPReAD) is a simple and versatile means of extending the concentration range of single molecule measurements into the cellular regime while minimizing potential artifacts and perturbations of protein complex stoichiometry.

**SIGNIFICANCE STATEMENT:** Quantifying the composition and stoichiometry of protein complexes in live cells is critical to understanding mechanisms involved in their function. Here we detail a method in which protein complexes are assembled intracellularly at physiological concentrations, but then diluted to levels suitable for single-molecule fluorescence observations while still within a cellular environment. The technique permits the use of common single molecule analysis techniques such as stepwise photobleaching quantification and fluorescence correlation spectroscopy to determine stoichiometry and functional interactions while avoiding artifacts that may occur from the use of detergent isolation methods or from the artificially low expression levels sometimes used to attain single molecule observation levels.

## INTRODUCTION

Dynamic networks of protein interactions underlie much of cell biology. A key goal of biomedical science is to understand the nature of these interactions and elucidate how they change in response to various extracellular cues. The native subunit stoichiometry of protein complexes often plays an important role in determining and regulating a protein’s function. Screening methods such as yeast-two hybrid analysis or phage display are useful for identifying potential binding partners in a high-throughput manner, but generally ignore biological context (1). Ensemble approaches that rely on co-immunoprecipitation or fluorescence spectroscopy can more accurately capture interactions within the cellular environment and used to examine changes that occur in response to external stimuli (1, 2). However, these bulk ensemble averaged measurements yield little information about the stoichiometry of subunits within complexes. Single molecule methods have the sensitivity to probe single protein complexes and quantitatively report on their individual architectures, further enabling the detection of critical subpopulations or heterogeneities.

Early uses of single-molecule fluorescence for subunit counting relied on artificially low expression levels in order to resolve individual protein complexes (3). However, non-physiological concentrations during assembly can shift binding equilibria and alter normal stoichiometry. More recently, a single-molecule pull-down (SiMPull) approach has been developed so that complexes can be assembled at native expression levels, extracted into a cell lysate, and then captured on an antibody-coated slide for single-molecule imaging (4). Antibody concentrations and lysate dilutions can be tuned to maintain single-molecule resolution without compromising intracellular assembly conditions. Although SiMPull has been used to successfully measure the subunit stoichiometry of membrane receptors, mitochondrial proteins, virus replication initiation, nuclear export complexes, and signaling complexes, the use of detergents for isolation and subsequent wash steps has been noted to affect the integrity of some macromolecular assemblies, particularly of membrane receptors, and therefore the physiological relevance of stoichiometry data (5).

Here, we introduce a simple detergent-free method to examine single protein complexes assembled at normal physiological concentrations in a near-native environment. Two cell populations—one containing a protein complex of interest and the other lacking it—are co-plated on a coverslip and fused into large syncytia. Protein diffusion within these syncytia results in a dilution of labelled complexes permitting their examination at reduced concentrations. Dilution factors are controllable by varying the co-plating ratio and can be made sufficiently high to resolve single membrane protein complexes in TIRF for stepwise photobleaching and brightness analysis, 2D membrane fluorescence correlation spectroscopy (FCS), two-color single molecule colocalization or single molecule FRET experiments. Cytosolic proteins are diluted as well and high-quality *in vivo* FCS and FCCS data can be obtained using the method. We call our approach Single Protein Recovery After Dilution (SPReAD), as it yields concentrations suitable for single molecule imaging after physiological oligomer assembly.

## RESULTS

### Formation of large syncytia using an inducible VSVG

A stable cell line with conditional expression of vesicular stomatitis virus G protein (VSVG) was created and used to initiate controlled cell fusion between the VSVG expressing cell line and cells expressing a labeled protein of interest. VSVG is a well-characterized fusogen that can be reversibly activated by a short pH drop (6). To dilute protein complexes for stoichiometric analysis, doxycycline-inducible VSVG-expressing U2OS cells were mixed with cells expressing the target protein typically at a 10:1 ratio (VSVG cells to target cells) and incubated at pH 5.5 for 5 minutes. After activating VSVG in a confluent monolayer of the mixed cell culture, we observed rapid formation (<1 hour) of massive syncytia and diffusion of labeled proteins producing a uniform distribution in which single molecules (adrenergic receptors in this case) were clearly visualized (Fig. 1a and b, right). The resulting images yielded punctate spots that resembled what we found using single molecule pull-downs of detergent-isolated protein from the same cells (Fig. 1b, left). These results suggest that substantial dilution factors may be attained in time intervals comparable to handling times for cell lysate preparation, implying that the two approaches have similar bounds for detecting transient, non-covalent oligomerization. However, SPReAD has the advantage of not requiring a detergent isolation step, which could disrupt non-covalent interactions, and the initial intracellular complex formation is carried out under normal cellular conditions before cell fusion.

**Figure 1.**
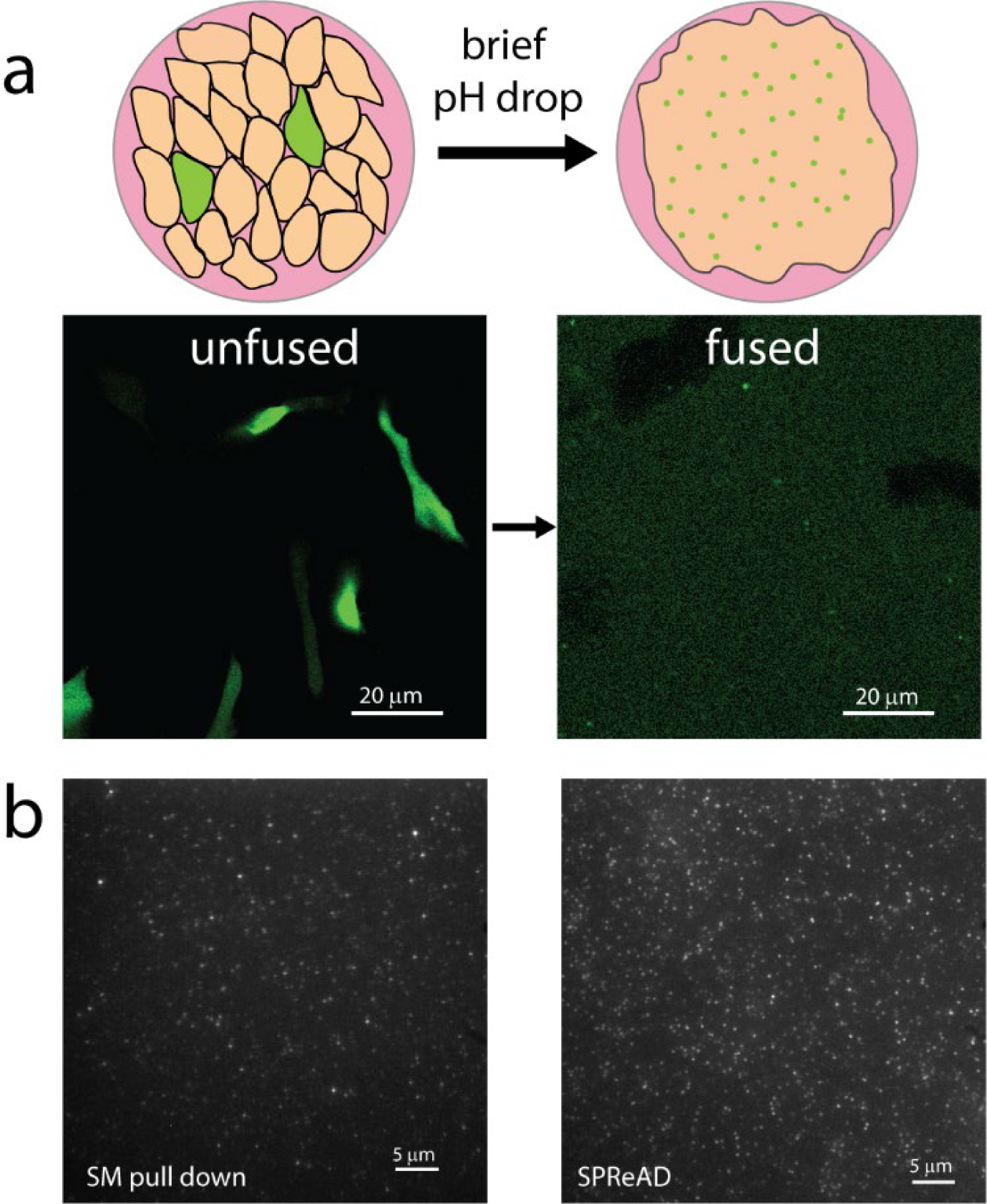
Use of Single Protein Recovery After Dilution (SPReAD) for single molecule imaging. (a) Cells expressing a labeled protein-of-interest (green) are co-plated with a stable U2OS cell line (orange) which express VSVG after doxycycline activation. A brief incubation in low pH (5.5-6.0) buffer initiates membrane fusion, after which protein complexes diffuse out of their parent cells into the larger syncytium. (b) mNG-β_2_AR protein complexes prepared for single molecule imaging by detergent isolation and biotin-streptavidin pull-down (left) and mNG-β_2_AR protein complexes in the syncytium membrane after VSVG-mediated fusion (right).

The dynamics of cell fusion and long-term viability of syncytia was visualized using time-lapse bright-field imaging (SI Appendix Fig S1a) which showed that fusion was accompanied by a loss of cell boundaries approximately 40 minutes after pH drop. For the next ∼4-5 hours, the syncytium remained bound to the coverslip and displayed few morphological changes. Thereafter, adhesion was slowly lost over the course of 12 hours and, at 20 hours, concerted cell death was observed. This suggests there is a 4-5 hour period during which cells are fused, but otherwise minimally perturbed. Cells may be imaged live during this window or fixed for later observation. Local spreading by diffusion of both cytosolic and membrane proteins occurs rapidly (SI Appendix Fig. S1b, c and d) and single molecule compatible levels are reached in 20-30 minutes.

Large-scale fusion was possible in all mammalian cell lines tested as expected due to the broad tropism of VSVG (SI Appendix, Fig. S2). This provides an additional means of experimental control by which background protein factors can be tuned by choice of cell type.

The formation of syncytia is a key step in the development of various mammalian tissues, including bone, muscle and placenta (7). In these cases, cell fusion is well regulated and part of the normal developmental program. Cell fusion can also play a role in the progression of disease. Many enveloped viruses trigger fusion between an infected cell and its neighbors, resulting in new and abnormal hybrids. Accidental cell fusion, both due to viral infection and otherwise, has also been implicated in cancer, where polyploid cells display high levels of chromosomal instability and may acquire tumorigenic phenotype (8). These natural examples of cell fusion suggest that large syncytia, at least during the first few hours after fusion, constitute a system in which oligomerization may be studied while preserving the biophysical environments of membrane and cytosolic protein complexes.

### Dilution of labelled cytosolic proteins by cell fusion improves *in vivo* Fluorescence Correlation Spectroscopy

Since its invention in the 1970s, fluorescence correlation spectroscopy (FCS) has become a valuable tool for investigation of molecular transport and interactions (9). Autocorrelation analysis provides information about diffusion, per-particle brightness and local concentrations, while two-color fluorescence cross-correlation spectroscopy (FCCS) can probe molecular associations (10). FCS and FCCS can be used inside living cells, but cellular proteins are typically expressed at intracellular concentrations outside of the working range for FCS studies. Furthermore, standard FCS and FCCS fitting models assume an infinite pool of diffusive species such that molecular motions are unconstrained and photobleaching is inconsequential. However, this is hardly the case within the cellular environment and these assumptions are known to lead to artifacts (11). Cell fusion is a promising means to address both of these limitations as concentrations can be tuned to fall inside the optimal range for FCS, and the relatively large size of the syncytium serves to alleviate the effects of constrained motion and photobleaching that can occur during live-cell FCS.

For cell fusion to function as a dilution strategy, protein complexes must be sufficiently mobile to diffuse out of their parent cells into the larger syncytium. Proteins confined to specific organelles or stably tethered to cytoskeletal components may fail to satisfy this criterion; however, many transcription factors and signaling complexes have mobile cytoplasmic fractions and are candidates for cell fusion-based dilution and single molecule analysis. The kinetics of syncytium formation and protein mobility determine the optimal timeframes for imaging and fixation after fusion is initiated. Time-lapse imaging revealed that membrane fusion was immediate and synchronized across the imaging dish, with cytosolic proteins beginning to escape their parent cells within 2 minutes of pH drop (SI Appendix Fig. S3). The initially heterogeneous fluorescence distribution was continually reshaped by diffusion until reaching a uniform steady-state level after ∼30 minutes. The equilibration time depends on the size of protein complexes being studied, their interactions with static cellular components, experimental conditions and the ratio of expressing and non-expressing cells. Overall, the kinetics of cell fusion and protein redistribution provide two possible modes of measurement. Measurements made in the non-equilibrium mode, prior to equilibration of the protein distribution, will most accurately report on the stoichiometry of weakly interacting protein complexes because assemblies have less time to dissociate before recording. However, concentration measurements at this stage will be heterogeneous across the imaging dish. In contrast, equilibrium mode measurements can be used to back-calculate the average intracellular concentrations prior to cell fusion (based on a known co-plating ratio), but may provide less accurate stoichiometric measurement of complexes with the fastest dissociation rates. Overall, this flexibility renders SPReAD as a versatile method for quantification of both oligomeric state and cellular expression levels.

To determine the range of dilutions possible, non-fluorescent VSVG-expressing cells were mixed with cells stably expressing mNeonGreen (mNG) at various co-plating ratios. After fusion, the fluorescence signal per unit area dropped in proportion to the co-plating ratio (SI Appendix, Fig. S2). Absolute numbers for syncytial concentrations were obtained by fluorescence correlation spectroscopy (FCS) and showed a similar trend, deviating only higher concentrations where FCS-based quantification is unreliable. We found that fusion-based dilution could be used to adjust cytoplasmic levels of an expressed protein over ∼two orders of magnitude. Importantly, larger dilutions brought cytosolic levels down to the sub-100 nM range, where the most quantitative and robust correlation spectroscopy measurements can be made.

To explore the benefits of using SPReAD for intracellular FCS measurements, we compared FCS data obtained from unfused cells with those from syncytia (Fig. 2a). In cells, transient mNG expression from a CMV promoter often failed to produce suitable autocorrelation curves, due to the high cytosolic concentration of labeled protein following transfection. FCS data is typically not useable when fluorophore levels exceed ∼1 μM, which is well within the range of normal intracellular protein concentrations. In practice, one either picks cells with low enough expression to obtain useable correlation curves or carries out whole cell photobleaching to reduce the fluorescent species concentration to FCS-compatible levels (12). Both of these options have clear biological drawbacks – either biasing the results by selecting only the low expressing cells, or phototoxicity from the bleaching method. We found that autocorrelations in unfused cells had an average dwell time of 2.2 ± 1.3 ms corresponding to a diffusion coefficient of 10 ± 5.8 μm^2^/s. We attribute the large deviations in the measured values (∼50%) from overall poorer data quality due to the measurements being made at the higher than ideal fluorophore concentrations, and to altered mobility near bounding membranes or organelles within the single cells. We often saw artifacts due to photobleaching, which manifest as a change in G(0) over time (SI Appendix Fig. S4). In comparison, dwell times and G(0) values from syncytial data FCS curves showed much less variation due to the larger homogenous pool of diffusing fluorophores. Syncytial FCS curves yielded dwell times and diffusion coefficients (1.2 ± 0.1 ms; 13 ± 1.1 μm^2^/s) similar to the unfused cells but with much less variation.

**Figure 2.**
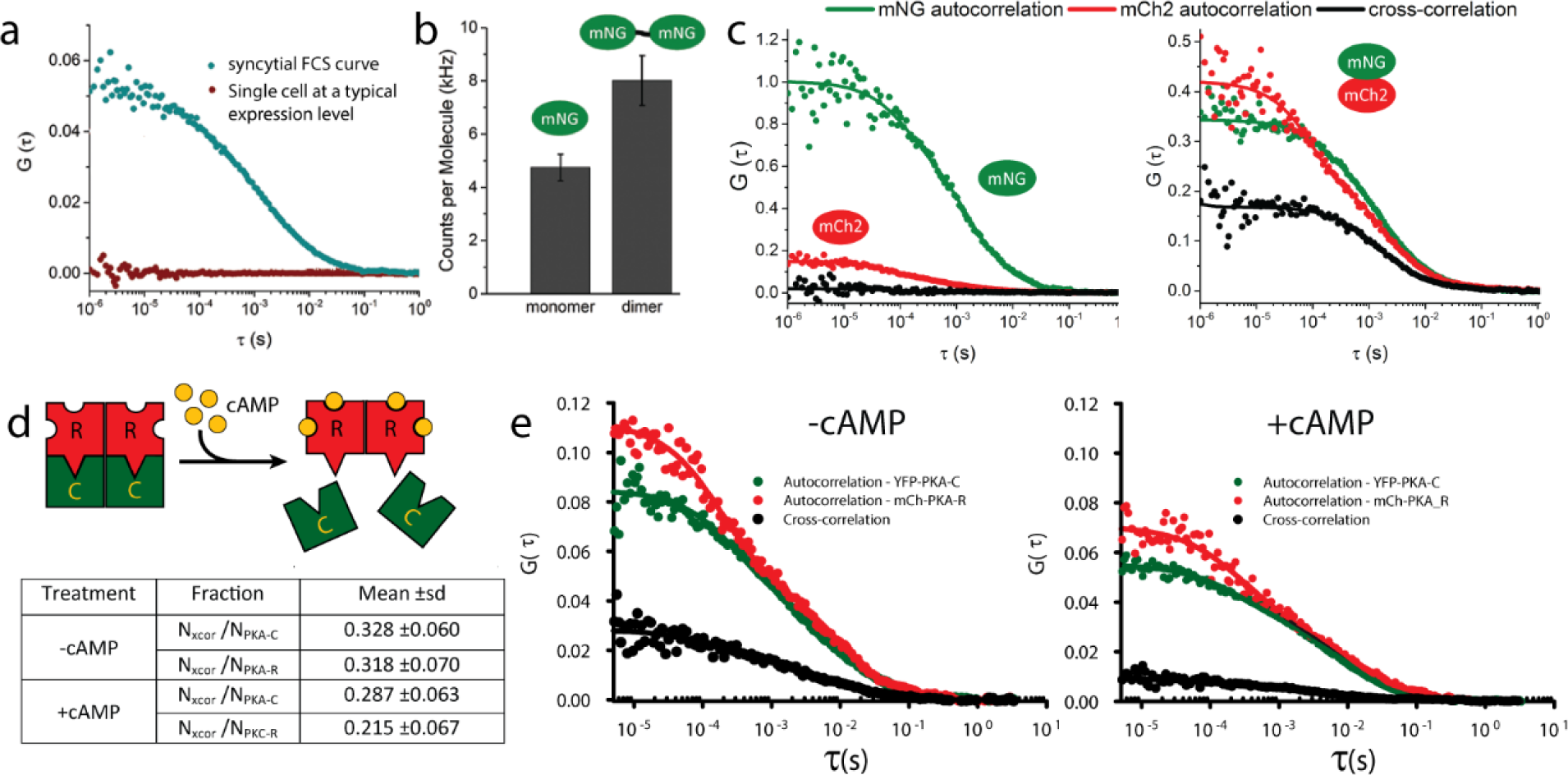
SPReAD improves in vivo fluorescence correlation spectroscopy (FCS) measurements. (a) FCS curves from syncytia are of uniform high quality since the concentrations can be set to FCS-compatible levels, compared to expression levels generally found in single transformed cells. (b) Brightness based on counts per molecule can be used to discriminate between monomeric and dimeric species in the cytoplasm of large syncytia and is useful for studying the stoichiometry of a single component within oligomers. (c) Cross-correlation spectroscopy is useful for studying heteromeric interactions. mNeonGreen and mCherry produce an appreciable cross-correlation (black line) when covalently joined (right) but not when co-transfected (left). In both cases, auto-correlations for each protein are clearly discernable. (d and e) FCCS in syncytia can be used to study functional differences in protein oligomerization. Protein Kinase A regulatory and catalytic subunits form complexes in the baseline state, repressing activity. Stimulation of adenylyl cyclase generates cAMP, causing PKA dissociation and increased activity. Table values in (d) represent data from 13 experiments.

Brightness analysis and two-color fluorescence cross-correlation spectroscopy are two valuable methods for studies of protein-protein interactions within the cellular environment (10, 13). To evaluate these techniques in conjunction with cell fusion, we compared measurements made with covalent dimers of fluorescent proteins to the corresponding monomeric proteins. mNG dimers were found to be 1.7 times brighter than monomers (Fig. 2b). Assuming minimal quenching effects, this suggests a maturation efficiency of 80-85% for mNG, which is on par with that of other green/yellow fluorescent proteins. Two-color fluorescence cross-correlation spectroscopy (FCCS) of mNeonGreen-mCherry2 covalent dimers yielded a 58% dimer population while a co-transfection of the monomeric proteins showed negligible cross-correlation amplitude (Fig. 2c). In addition to brightness and cross-correlation analyses, other methods such as photon counting histograms, dwell time distributions, photon anti-bunching and single-molecule FRET have been used to examine oligomerization states and could be aided by SPReAD sample preparation.

Next, we used syncytial two-color FCCS to study the oligomerization of protein kinase A (PKA), a Ser/Thr kinase that functions in the cAMP-dependent pathway of G-protein coupler receptor (GPCR) signaling. Upon GPCR activation, adenylyl cyclase catalyzes the conversion of ATP into cAMP, causing protein kinase A (PKA) regulatory subunits to dissociate from catalytic subunits, which are then free to phosphorylate downstream targets. Syncytial FCS of YFP-tagged catalytic subunits and mCherry-tagged regulatory subunits revealed a substantial cross-correlation, indicating functional repression in the baseline state (Fig. 2d, e). Upon stimulation with the adenylyl cyclase activator forskolin and the phosphodiesterase inhibitor IBMX, cross-correlation amplitudes decreased, reflecting cAMP-induced dissociation of subunits, and mirroring previous efforts using FCS in live cells or SiMPull with cell lysates (10, 11, 14). SPReAD increases the usefulness and robustness of FCS and FCCS for cell based measurements by allowing for target complex formation at more physiological intracellular concentrations and by mitigating complicating effects from confined cellular volumes.

### Single molecule imaging of membrane protein complexes

Most membrane proteins are freely mobile in two dimensions, unless tethered to intracellular actin. Membrane residing protein complexes are of significant interest to biomedical research, representing 23% of all ORFs in the human genome and being the target of >60% of pharmaceutical drugs (15). The biomedical significance of membrane receptors has motivated comprehensive investigation of their basic structures and mechanisms of action. Oligomerization is known to play a role in the function of many major receptor types (metabotropic, ionotropic and tyrosine kinases) and thus, considerable effort has been made to elucidate their interaction profiles. From a single-molecule perspective, subunit counting in oocytes has been the widely used approach, with many receptors being studied after controlled mRNA injection to limit receptor levels (16). However, the concentration-dependence of oligomerization may be at odds with the sub-physiological expression levels employed in this technique and cell type specific post-translational modifications occurring in the Golgi and ER required for native oligomer formation may be lacking (17). We show that cell fusion combined with single molecule imaging lifts this restriction and allows single molecule imaging after physiological assembly of receptor complexes in a cell type required by any specific biological constraints.

We undertook a series of experiments designed to demonstrate the unique utility of SPReAD for single molecule imaged based measurements of membrane protein stoichiometry. In these experiments, we examined differences between the results obtained by SPReAD preparations and single molecule pulldowns to determine if detergent isolations had a notable effect, as well as investigations in the oligomeric state of several well-studied membrane protein complexes.

### Beta-2 adrenergic receptor stoichiometry

The adrenergic receptors (ADRβ1-3) are class A G protein coupled receptors (GPCRs) that are targets of catecholamines such as adrenaline and noradrenaline. Oligomerization of class A GPCRs is still controversial, with reports of ADRβ1 and ADRβ2 forming homodimers and heterodimers (18); while more recent reports disputing this finding (19). We used mNG-tagged ADRβ2 expressed in U2OS cells to assess dimer formation and compare our method to single molecule pulldown results. mNG-ADRβ2 expressing cells were co-plated with VSVG-expressing neighbors to dilute membrane receptors from the initial high expression levels. After cell fusion and incubation at 37°C for 1 hour, individual receptor complexes were clearly discernible and mobile within the plasma membrane, displaying similar kinetics to measurements made in living cells (Fig. 1b, SI Appendix Video 1). Single particle tracking of mNG-ADRβ2 (SI Appendix, Fig. S5) also confirmed this observation. The receptor concentration distribution was more heterogeneous across the syncytium than we saw with cytosolic proteins even ∼1 hour after fusion due to the slower diffusion rate for proteins in the membrane compared to cytoplasm. However, there were still numerous fields of view with uniform sparse distributions ideal for single molecule imaging (Fig. 1b, right panel).

Syncytia were fixed with paraformaldehyde to immobilize receptor complexes and facilitate stoichiometry determination by stepwise photobleaching and two-color single molecule co-localization methods. mNG-ADRβ2 puncta showed distinct bleach steps (Fig. 3a, SI Appendix Fig. S6, top row). Analysis of the receptor population revealed that ADRβ2 was evenly distributed between monomeric and dimeric states, with 25% of photobleaching traces showing two bleach steps (Fig. 3a, fourth bar group), signifying a 36% dimer population after accounting for mNG’s maturation efficiency (Supplementary Methods). In order to compare the effects of n-Dodecyl β-D-maltoside (DDM), a detergent commonly used in cell lysis for pulldown experiments we carried out single-molecule pull-down (SiMPull protocol) experiments on the same mNG-ADRβ2 expressing cells and found that dimer fractions differed from what was observed with SPReAD (Fig. 3a, first three bar groups). Using a standard detergent isolation and single molecule pulldown procedure the dimer fraction was never higher than 10%, indicating that in some cases detergent isolation methods can introduce significant errors in single molecule stoichiometry determinations.

**Figure 3.**
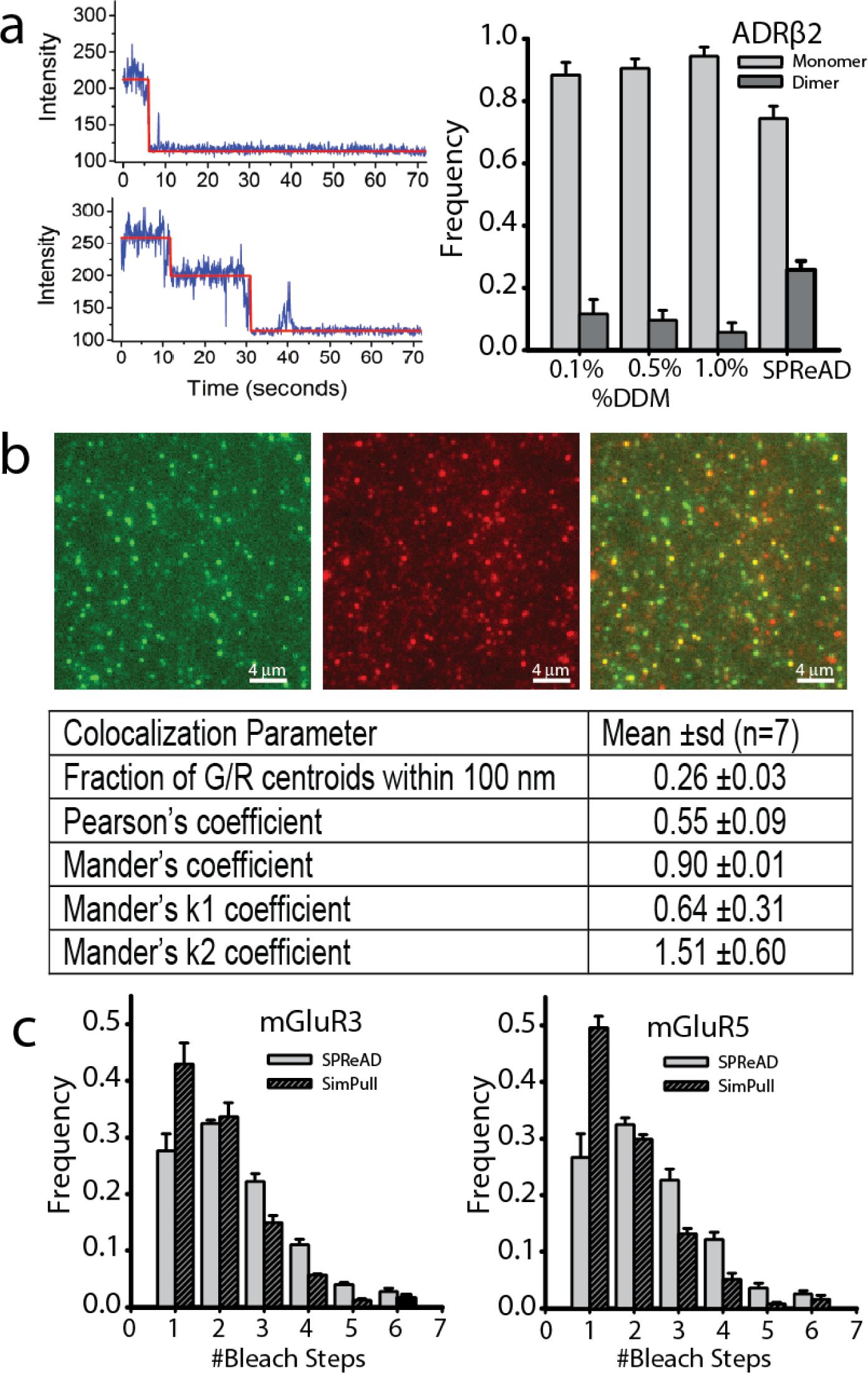
Single Protein Recovery After Dilution (SPReAD) for single molecule imaging avoids potential artifacts of detergent isolation. (a) Typical monomer (top-left) and dimer (bottom-left) mNG-ARβ2 bleach step traces obtained from the SPReAD-prepared samples. Right: The elimination of detergent isolation artifacts was demonstrated by measuring the mNG-ARβ2 dimer to monomer ratio from pull down experiments using mNG-ARβ2 isolated at three different detergent concentrations (first three groups of bars) and from SPReAD preparations (fourth bar group). Data is the mean ± SEM, n = 3. These experiments showed a significantly larger fraction of mNG-ARβ2 dimers using SPReAD, which we take to be a more accurate measure of the physiological dimer ratio. (b) Two-color single molecule colocalization using SPReAD. Top: Green, red and overlay images of mNG-ADRβ2 and mCherry-ADRβ1 revealing colocalization of adrenergic receptor subunits after cell fusion based on PSF colocalization analysis. Bottom: Averaged colocalization results from single molecule centroid based colocalization analysis and from conventional image-based colocalization methods (n = 7). (c) Comparison of metabotropic glutamate receptor (mGluR3 and mGluR5 – both known to form homodimers) complex stoichiometry in samples prepared via SPReAD, or via lysis for SimPull (mean ± SEM, n = 3). We demonstrate that detergent-isolated single molecule pull down experiments show a larger monomer fraction relative to SPReAD, which we interpret as an effect of the isolation treatment or subsequent wash steps on complex stability.

### Heteromeric complex stoichiometry measurements of ADRβ2-ADRβ1 in fused cells

To demonstrate the ability to probe heteromeric associations with cell fusion and single molecule imaging, we co-expressed mNG-ADRβ2 and mCherry-ADRβ1 in U2OS cells. After fusion with VSVG-expressing cells both color channels showed distinct puncta in the syncytia and the overlaid image clearly displayed overlapping spots (Fig. 3b). The degree of colocalization was quantified by the standard pixel level methods (Pearson’s and Mander’s coefficients) and at the single molecule level by PSF fitting and determining the fraction of spots with a nearest neighbor in the opposite color within 100 nm (Table in Fig. 3b). After fusion, the respective color channels showed a high degree of colocalization with 26% of mNG-ADRβ2 spots overlapping with mCherry-ADRβ2 spots as determined by centroid localization (Fig. 5b), which predicts a 40-50% colocalization, when corrected for missed colocalized pairs from non-fluorescent proteins. The lower brightness of mCherry prohibited accurate stepwise photobleaching measurements in the red channel; however, based on the single molecule colocalization observed and statistical analysis of heterodimer formation we conclude that the affinity for ADRβ2 – ADRβ1 heterodimer formation may be about equal to that of homodimer formation.

### Metabotropic glutamate receptor stoichiometry

The metabotropic glutamate receptors mGluR3 and mGluR5 are known to function as covalently bound homodimers via a cysteine bridge assembled in the ER prior to membrane trafficking (20). We generated stable HEK293T cell lines constitutively expressing a mNeonGreen labeled metabotropic glutamate receptor, either mNG-mGluR3 or mNG-mGluR5 and co-plated them with VSVG-expressing U2OS cells. Fusion was induced and syncytia were formed and fixed with paraformaldehyde as described above. After fusion, individual receptor complexes were able to be resolved in many areas of the dish, demonstrating fusion between different cell types with proteins able to diffuse throughout the heterogeneous membrane of these syncytia.

Stepwise photobleaching experiments performed on the syncytia found that 33.5 ± 0.5% and 33.8 ± 0.8% of the traces showed two bleach steps for mNG-mGluR3 and mGluR5 respectively, confirming that complexes formed prior to membrane trafficking were preserved during the fusion process (Fig. 3c). Single-molecule pull-down experiments were performed on the same cell lines using 1% DDM in the cell lysis buffer. In contrast to what we noted earlier for adrenergic receptors, we found only slightly lower dimer fractions for both mGluRs (29.9 ± 0.8% for mGluR3 and 28.1 ± 0.4% for GluR5 - Fig. 3c), possibly due to disulfide linkages between mGluR monomers. Heterodimeric mGluRs have been proposed and single molecule methods such as SPReAD would be a useful means of detecting them. Heterodimeric mGluRs have important implications since the pharmacology of heterodimers may differ significantly from the better characterized mGluR homodimers (21).

### Detection of higher-order oligomeric membrane protein complexes – CRAC channel subunit Orai1 stoichiometry

We also examined the subunit stoichiometry of Orai1, a calcium-selective ion channel that forms the central pore of the calcium release-activated channel. The functional stoichiometry is currently unresolved, with claims of either a tetrameric or a hexameric configuration (22–24). Using SPReAD we found that most Orai1 puncta bleached in 1-6 steps (Fig. 4a), which supports the hexameric model proposed in (22), especially after correction of the raw bleach step distribution for an estimated dark fraction of mNeonGreen is taken into account. Assuming a 20% successful protein folding rate (Supplementary Methods), the corrected distribution has a weighted mean of five Orai1 subunits per complex. Although not fully hexameric on average, this result may be due to the interference by the mNG moiety and/or the presence of unlabeled Orai1 in mammalian cells. Targeted knockdown of endogenous proteins or careful choice of cell lines may be used to refine understanding of physiologically relevant oligomerization in specific tissue types.

**Figure 4.**
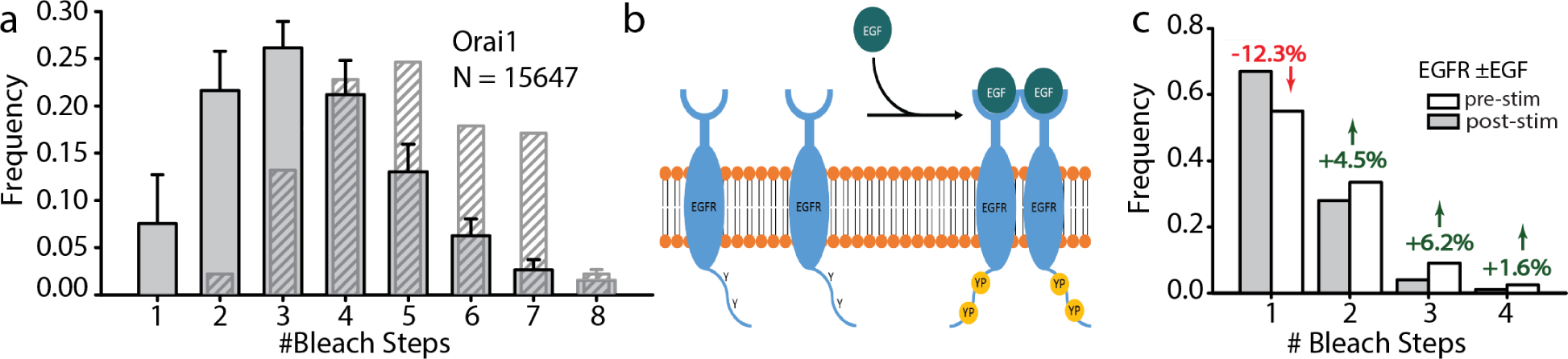
Application of SPReAD to detect higher-order oligomer membrane protein complexes and measure stoichiometric changes due to ligand binding. (a) The functional form of the CRAC channel continues to be a contested issue, with claims that Orai1 subunits adopt either a tetrameric or a hexameric configuration. The raw bleach step data from 15647 fluorescent spots analyzed from SPReAD syncytia made from Orai1 expressing cells yielded a weighted mean step number of 3.5 (gray bars). Hatched bars represent an estimate of the actual subunit fractions assuming a 0.8 fluorescent fraction for mNG and indicate a weighted mean subunit number of 5.0. (b) Epidermal growth factor binding stimulates EGFR dimerization and autophosphorylation of tyrosine residues on EGFR’s cytoplasmic tail, leading to recruitment of downstream signaling proteins. (c) SPReAD bleach step histograms of EGFR oligomerization before (gray bars) and after (white bars) EGF stimulation. Although EGFR is largely monomeric prior to growth factor addition, there is a substantial dimer fraction as well. After stimulation, the dimer and higher-order oligomers fractions increase, while the monomer fraction drops.

### Ligand-dependent oligomerization of epidermal growth factor receptor

One of the primary evolutionary advantages conferred by oligomerization is the development of new modes of regulatory control of protein activity. Allosteric oligomerization is known to play a role in the mechanisms of both metabotropic receptors and receptor tyrosine kinases, with extracellular ligands modulating the formation of dimers or higher order structures. Monomer-oligomer transitions can prime receptors for downstream signaling events, such as post-translational modifications or the recruitment of adaptor proteins. As an example of the use of SPReAD to detect ligand-dependent multimerization, we looked at epidermal growth factor receptor—a member of the receptor tyrosine kinase family whose abnormal regulation has been implicated in a number of human cancers (25). The canonical model for receptor activation asserts that EGFR is monomeric in the plasma membrane prior to stimulation, whereupon it is driven to dimerize upon binding of its cognate ligands, resulting in autophosphorylation of tyrosine residues on its cytoplasmic tail and recruitment of specific effector proteins (Fig. 4b).

Although the EGFR pathway has been extensively studied using both bulk and single molecule approaches, there are still open questions about receptor oligomerization. There is increasing evidence that pre-formed dimers of EGFR exist on the cell surface prior to ligand stimulation and that EGFR is capable of forming higher-order oligomers that may function in receptor activation (26–28). To examine each of these possibilities, we expressed a mNeonGreen-tagged EGFR (mNG-EGFR) on the cell surface and performed stepwise photobleaching measurements in fixed syncytia. Even in its baseline state, EGFR was found to be significantly dimeric, with 29% of traces bleaching in two steps (Fig. 4c). Upon stimulation with EGF, this dimer fraction increased substantially and some higher-order oligomers (trimers and tetramers) were observed. Together, these results support a model where at least some EGFR signaling is accomplished by conformational changes in pre-formed dimers and trans-activation by higher-order oligomers. The use of SPReAD to study ligand-dependent oligomerization of EGFR validates its potential for studying transient interactions.

## DISCUSSION

By achieving detergent-free dilution of protein complexes after physiological assembly, SPReAD facilitates measurements of subunit stoichiometry for both cytosolic and membrane-bound oligomers. Furthermore, the use of VSVG as a means of accomplishing cell fusion is highly efficient and requires only a simple buffer exchange. In contrast, existing methods for probing oligomerization are significantly more complex or disruptive. Use of stimulated emission depletion to reduce excitation volumes by >100-fold can extend the upper limit on FCS measurements (29) but requires complicated optics and increases photobleaching and phototoxicity. As mentioned earlier, single molecule pull down approaches can probe molecular heterogeneity in oligomerization but require extraction of protein complexes from their native environment (4, 30). Meanwhile, efforts to apply conventional imaging or localization microscopy to stoichiometry analysis rely on *a priori* assumptions about protein distribution or fluorophore blinking (31, 32). Compared against these other methods, SPReAD offers unique advantages, affording single molecule sensitivity for oligomerization studies while maintaining a more physiologically relevant setting.

Although the use of cell fusion for dilution is both simple and rapid, the dynamics of various intracellular processes need to be considered when interpreting results. Syncytia form almost instantly after pH drop, but protein redistribution is diffusion-limited and thus, much slower for membrane-bound proteins undergoing 2D diffusion compared to cytosolic proteins moving much more rapidly in 3D. This yields two possible modes of analysis: an equilibrium mode, where the final concentration of labeled protein complexes is uniform and proportional to the initial concentration (divided by the co-plating ratio), and a non-equilibrium mode, where concentrations across the imaging dish are non-uniform. The latter mode, typically carried out 10-45 minutes after buffer exchange for the proteins studied here, is most conducive to measuring subunit stoichiometry as it minimizes time during which non-covalent protein complexes can dissociate. Beyond these considerations of diffusion and dissociation, syncytia appeared to be morphologically stable for 5-6 hours but it is still largely unclear how the intracellular environment is reshaped during the fusion process. Understanding how syncytium formation affects major signaling pathways will be critical for proper interpretation of SPReAD results.

A concern with our approach may be that the use of a brief pH drop to initiate VSVG-mediated cell fusion may effect complex stoichiometry. We note that in our comparisons between SPReAD and single molecule pulldowns, we consistently saw higher levels of oligomerization using our cell fusion method compared to detergent isolated preparations, indicating that the pH jump is at least less disruptive than detergent pulldown for the proteins we studied. Furthermore, despite the extracellular pH drop, we found that intracellular pH is largely unchanged (SI Appendix Fig. 8), and therefore proteins and protein complexes en route to the membrane are likely unaffected. Proteins trafficked to the membrane during the fusion time course occurring after the pH jump would be able to associate normally for complexes that oligomerize on the membrane (e.g. EGFR). It should also be possible to achieve in situ protein dilutions similar to what we show here using alternative viral fusogens, such as Reovirus Fast proteins (33), which do not require a pH jump to activate.

A number of strategies may be used to augment the SPReAD technique and build upon its versatility. For accurate measurement of physiologically relevant interactions, endogenous proteins can be labeled using prevalent genome editing techniques or, ideally, primary cells can be extracted from genetically modified organisms to understand tissue-specific phenotypic variation. Future work may also extend SPReAD applications beyond the cytoplasm and plasma membrane by making use of membrane contact sites between organelles, examining proteins that exchange between the cytoplasm and other compartments or by retargeting of proteins through signal sequence engineering.

By removing limits on expression levels compatible with single molecule experiments without requiring chemical agents for dilution, SPReAD permits minimally perturbative measurements in a variety of cell lines. Aside from the FCS- and stepwise photobleaching-based analyses of subunit stoichiometry highlighted here, we expect SPReAD to enhance other methods traditionally limited to working at low concentrations such as smFRET, single-particle tracking and single molecule spectroscopy, thus providing a powerful addition to the single molecule toolkit.

## METHODS

### Cloning of inducible VSVG and labelled proteins

To avoid the deleterious effects of long-term VSVG expression, the coding sequence for VSVG (Addgene #8454) was cloned into the BamHI and EcoRI sites of the lentiviral pLV Puro Tet vector for doxycycline-inducible expression. A constitutively expressed mNeonGreen lentiviral plasmid was produced by excising mNeonGreen from mNeonGreen-N1 (Allele Biotech) using NheI and NotI and subcloning into pCDH-puro (System Biosciences). This resulting plasmid has been deposited with Addgene (plasmid #82724).

Synthetic dimers of fluorescent proteins were produced by placing a helical linker A(EAAAK)_5_A after the mNG sequence in mNG-C1 (between the BspEI and BglII sites). mNG or mCh2 were then PCR amplified and placed after this linker (between NotI and SpeI sites) to generate mNG-mNG or mNG-mCh2, respectively. The rigid helical linker spaces the fluorescent protein domains further apart to reduce energy transfer (34, 35). pCDH-puro and mNG-C1 were both digested with NheI and BamHI to excise the fluorescent protein and place it into the pCDH lentiviral plasmid to generate pCDH-puro-mNG, which was used to produce a stable mNG cell line.

mNG-tagged ADRβ2 and EGFR were generating by cloning into the pSNAPf-ADRβ2 backbone (New England Biolabs). mNG was amplified by PCR from mNG-C1 and placed between the EcoRI and SbfI sites of pSNAPf-ADRβ2 (replacing the SNAP tag) to yield mNG-ADRβ2. Site-directed mutagenesis was used to remove a ClaI site from wildtype EGFR. This mutated EGFR was then PCR amplified and placed between the SbfI and XhoI sites of the pSNAPf-ADRβ2 plasmid, replacing ADRβ2. The EGFR signal sequence was purchased as a gBlock (Integrated DNA Technologies) and placed between the ClaI and BmtI sites to generate mNG-EGFR. Lentiviral versions of mNG-ADRβ2, mNG-EGFR, mNG-mGluR3, and mNG-mGluR5 were produced by amplifying each plasmid via PCR and digesting with XbaI and NotI to place the fusion protein after the CMV promoter in pCDH-puro. To make Orai1-mNG, Orai1-YFP (Addgene #19756) and mNG-N1 were digested with AgeI and NotI to remove YFP and replace it with mNG.

### Cell culture and generation of stable cell lines

U2OS human osteosarcoma cells were cultured in DMEM without phenol red, supplemented with 10% fetal bovine serum (FBS), sodium pyruvate, 1x GlutaMax and 1x antibiotic-antimycotic; all cell culture media and supplements were purchased from Life Technologies. For stable expression of VSVG under tetracycline control, U2OS cells were first stably transduced with the rtTA NeoR plasmid for the reverse tetracycline-controlled transactivator (rtTA) protein. Lentiviral particles were generated in HEK293 cells and used to transduce U2OS cells as previously described^32^. Stably transduced cells were selected using 700 μg/mL G418. U2OS rtTA cells were then transduced with pLV puro Tet-VSVG and selected using 2 μg/mL puromycin. Doxycycline was withheld from cell culture media until 24 hours prior to cell fusion. Stable mNeonGreen cell lines were produced by transducing U2OS Tet-VSVG cells with pCDH-puro-mNG-C1, pCDH-puro-mNG-ADRβ2 and pCDH-puro-mNG-EGFR and selecting with 2 μg/mL puromycin. Stable mNeonGreen-mGluR expressing HEK293T cell lines were produced by transducing cells with either pCDH-puro-mNG-mGluR3 or pCDH-puro-mNG-mGluR5 and enriched by fluorescence-activated cell sorting (BD Biosciences FACSAria).

### Substrate Preparation

To minimize glass autofluorescence and maximize cell attachment, plain glass-bottom dishes were cleaned and coated with fibronectin. Dishes were etched with 1 M KOH for 20 minutes, followed by a water and then PBS rinse. For fibronectin coating, dishes were incubated in 4% (3-Mercaptopropyl)trimethoxysilane (Sigma-Aldrich) in ethanol for 30 minutes, rinsed with ethanol, incubated with (N-γ-maleimidobutyryl-oxysuccinimide ester) crosslinker (4 mM in ethanol, Thermo Scientific), rinsed with ethanol and dried thoroughly in a sterile biosafety cabinet. Dishes were then incubated with 5 μg/mL fibronectin for 2 hours at room temperature, followed by overnight at 4°C, then rinsed with PBS and stored in PBS at 4°C until use (up to several weeks).

### Fusion Assay

U2OS Tet-VSVG cells were plated onto collagen coated glass-bottom dishes. After reaching confluence, fresh media with 2 μg/mL doxycycline was added and the cells were returned to a CO_2_ incubator for 24 hours. Cells were then fused by removing culture media, washing with PBS and incubating in fusion buffer (PBS with 25 mM MES, pH 5.5) for 1 to 5 minutes. Cells were washed with PBS and culture media was restored before returning cells to the CO_2_ incubator. Cell membranes and nuclei were labelled at various time points by incubating with 5 μg/mL Wheat Germ Agglutinin Alexa 647 (Life Technologies) and 5 μg/mL Hoechst in Hank’s balanced salt solution for 10 minutes prior to fixation with 4% paraformaldehyde. Fixed cells were imaged on a spinning disk confocal microscope (Olympus) with air objectives (40x/0.9, 20x/0.7 and 10x/0.4) and examined for syncytia formation.

### Confocal Microscopy and Fluorescence Correlation Spectroscopy

U2OS Tet-VSVG cells were transfected with FP control plasmids or FP-tagged PKA-subunits using Lipofectamine 3000; for cytoplasmic mNG measurements, stable U2OS mNG cells were used to accurately control the number of expressing cells. Serum-free Fluorobrite DMEM (Life Technologies) was used to minimize cellular autofluorescence. The two were mixed at various ratios and 5×10^5^ cells were plated in the well of a 14 mm diameter glass-bottom dish (collagen/fibronectin-coated) using doxycycline-supplemented media (2 μg/mL); additional media was added 2-12 hours after plating, after cells were visibly attached and spread. Cells were imaged on a confocal microscope (Zeiss LSM880). Fluorescence correlation spectroscopy was performed on the same instrument using the LSM880 32-channel GaAsP detector in photon counting mode. Standard FCS fitting equation were used (10) and further details of FCS data analysis is detailed in the SI Appendix. For non-PKA FCS measurements, the data was fit to a single component diffusion with triplet model. Absolute concentrations for cytoplasmic mNeonGreen were obtained by calibrating the focal volume with known concentrations of Alexa 488. From the two-color cross-correlation measurements, the average number of particles was determined using:

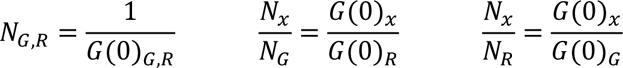

where N_G,R_ is the number of green or red particles, and N_X_/N_G_ and N_X_/N_R_ are the heterodimer fractions. For Protein Kinase A experiments, PKA-transfected U2OS cells were mixed 1:10 with non-expressing VSVG cells and incubated in doxycycline-supplemented Fluorobrite DMEM for 24 hours. Cells were then fused by a 5-minute incubation in fusion buffer and FCS was performed in syncytia one hour later. In order to maintain the same syncytial position for post-stimulation measurements, 2x cAMP-stim buffer (50 μM forskolin, 200 μM IBMX in Fluorobrite DMEM) was added directly to the imaging dish in equal volume to the residual media and a second FCS recording was initiated 5 minutes later. PKA data was fit to a two-component diffusion model (10)

### Single-Molecule Imaging After Cell Fusion

U2OS Tet-VSVG cells were transfected with FP-tagged receptor constructs and plated onto glass-bottom dishes with non-transfected cells at a ratio of 10:1 (non-transfected:transfected), as described above. After 24h of doxycycline induction, cells were fused and imaged live (1-2 hours after fusion) or fixed for stoichiometry/colocalization analysis. Cells were fixed with 4% paraformaldehyde for 3 hours in the dark at room temperature to eliminate residual mobility of membrane proteins after short fixation^33^. For mNG-EGFR experiments, the syncytia were stimulated with 200 ng/mL EGF 75 minutes after cell fusion was initiated and fixed 5 minutes later or fixed without EGF treatment.

### TIRF Microscopy

A custom-built azimuthal-scanning objective-TIRF microscope was used for single molecule imaging. Excitation at 488 nm and 561 nm were used to excite mNeonGreen and mCherry, respectively, and were directed to the sample using a quad polychroic (ZT405/488/561/640rpc, Chroma Technology) housed in the filter wheel. A beam telescope and focusing lens were used to create a collimated beam out of the objective (Olympus UApoN 100x/1.49), while a pair of XY galvanometer mirrors (Model 3210H, Cambridge Technology) controlled the angle of incidence. For the detection path, a TuCam adaptor (Andor) equipped with band pass filters (ET525/50 for mNeonGreen and ET605/52m for tdTomato) was used to split emissions onto two EMCCDs (Andor iXon 887 and 897 Ultra). Image coregistration was accomplished by acquiring brightfield images of a calibration objective (Zeiss LSM calibration objective) prior to each imaging experiment and ensuring that the images were coregistered to better than one pixel over the camera field-of-view through alignment of the detection pathway. Live-cell data was acquired at 37°C using an objective heater (Bioptechs), while fixed cell experiments were performed at room temperature. Coverslips were scanned for regions with a suitable density of molecules for single molecule analysis; regions with unfused fluorescent cells or too few/too many molecules were avoided. For bleach step analysis, 2000 frames were recorded at 10-30 Hz; laser intensity was kept low to mitigate blinking artifacts. For colocalization analysis, 20 frames were acquired and averaged during post-processing.

### Single Molecule Data Analysis

Photobleaching movies were analyzed using three different methods. Two are based on the use of a custom lab software package (ImageC.exe, written in C/C++ under Microsoft Visual Studio 2017) and the third was the use of a published software called Progressive Image Filtering (PIF) which is an automated software package written in Matlab (36). In both programs, molecules (PSFs) were first located automatically found by successive processing of the summed image stack to locate fluorescent puncta above a certain threshold that meets a specified Gaussian fit criterion. For each molecule, an ROI (typically 5×5) centered on the pixel containing the PSF centroid was created and the ROI mean values vs. time (frame) extracted from the stack. The ROI center pixel coordinate was readjusted slightly as needed as the data is extracted from the frames so that the brightest pixel is always at the center. ROI fluorescence traces of all the spots located are stored within the program and displayed as time trace plots for manual (i.e. visual) step counting in ImageC, or used for with the automated step-finding algorithms in ImageC or PIF. Both algorithms count the number of bleach steps based on signal noise and a user-set change in the trace count level that determines a valid step. Traces without discernible bleach steps were discarded. At least 700 molecules were analyzed for each sample. Further information on the programs used is provided in the Supplementary Methods section and in SI Appendix Fig S7.

For colocalization analysis, data from two EMCCDs were analyzed to find spots in both the green and red channels using either a custom MATLAB script or function built into our lab’s custom analysis program (ImageC). The PSFs were fit to a Gaussian model to determine center locations. A colocalization fraction was calculated to be the fraction of mNeonGreen spots with an mCherry spot less than 100 nm away.

## ACKNOWLEDGEMENTS

We would like to thank Yuri Lazebnik and Gary Whittaker for helpful discussions regarding VSVG-mediated cell fusion, Dave Holowka and Barbara Baird for EGFR plasmids and insight concerning cell-signaling pathways, Taekjip Ha and Yang Kevin Xiang for adrenergic receptor and protein kinase A plasmids, Matt Paszek for tetracycline-inducible plasmids and Julian Palacios-Goerger for assistance with lab-built incubator microscope imaging shown in the supplementary data section. A portion of the imaging data was acquired through the Cornell University Biotechnology Resource Center, with NIH (S10OD018516) funding for the shared Zeiss LSM880 confocal/multiphoton microscope and NSF (NSF-1428922) funding for the shared Zeiss Elyra super resolution microscope. This work was supported by NIH R33-CA193043 and R01DA030329 to WRZ, and NIH R01-GM101023 to PK and WRZ.

## AUTHOR CONTRIBUTIONS

W.R.Z. supervised the project. W.R.Z. and P.J.K. conceived the idea. A.S., A.VS. and M.S. assessed viability of cell fusion as a dilution strategy. A.S. built the microscope for single molecule imaging. A.S. and A.VS. generated the constructs and cell lines. A.S., A.VS, M.S. and A. Song conducted single molecule imaging experiments and data analysis; A.S. and W.R.Z. performed correlation spectroscopy; A.S., A.VS, W.R.Z. and P.J.K. wrote the paper with input from all other co-authors.

## SUPPLEMENTAL METHODS

### Time course imaging of cell fusion dynamics

The kinetics of cell fusion and subsequent protein diffusion were thoroughly investigated to guide the timing of single molecule experiments. These dynamics will vary for each protein of interest and thus, similar experiments may need to be conducted prior to FCS or stepwise photobleaching measurements. As an example of cytoplasmic proteins, U2OS cells expressing mNG were co-plated with non-expressing VSVG cells at a ratio of 1:10 or 1:100. Cells were fused using a 30-second incubation in fusion buffer and then returned to imaging media (Fluorobrite DMEM). The imaging dish was loaded onto a confocal microscope (Zeiss LSM 880) and a time-series was started 2 minutes after fusion was initiated. Both the 1:10 and 1:100 co-plating ratios were imaged at room temperature (23°C), while the former was also imaged at 37°C to examine the effects of temperature on protein diffusion.

Diffusion of mNG-ADRβ2 was investigated by fixing syncytia at various timepoints after cell fusion. Fusion was accomplished by a 30-second incubation in fusion buffer and cells were then returned to imaging media (Fluorobrite DMEM) at 37°C. Cells were fixed with 4% PFA for 3 hours, then washed with PBS, and imaged on a commercial TIRF microscope (Zeiss Elyra).

A custom-built brightfield microscope housed in a CO_2_ incubator was used to examine the time-course of cell fusion. The microscope consists of an x-y stage (Applied Scientific Instrumentation), ring illuminator and camera (Chameleon, Point Grey Research). Custom software allows for autofocus and tiling of the culture dish.

U2OS Tet-VSVG cells were plated and induced with doxycycline for 24h as described earlier. Cells were fused by a brief 30-second incubation in fusion buffer, washed with PBS and restored to normal culture media. Immediately, the dish was loaded into the incubator microscope and imaged for ∼30 hours.

### FCS/FCCS data analysis

For non-PKA experiments, data was fit to a single component diffusion with triplet model:

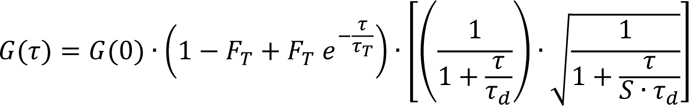

where τ_T_ and F_T_ are the triplet time and fraction, respectively, τ_d_ is the diffusion time, S is the structure factor for the focal volume and G(0) is the correlation at τ = 0. The structure factor was set to 10 for all fits. From the two-color cross-correlation measurements, the average number of particles was determined using:

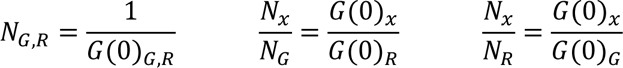

where N_G,R_ is the number of green or red particles, and N_X_/N_G_ and N_X_/N_R_ are the heterodimer fractions. Absolute concentrations for cytoplasmic mNeonGreen were obtained by calibrating the focal volume with known concentrations of Alexa488.

For Protein Kinase A experiments, PKA-transfected U2OS cells were mixed 1:10 with non-expressing VSVG cells and incubated in doxycycline-supplemented Fluorobrite DMEM for 24 hours. Cells were then fused by a 5-minute incubation in fusion buffer and FCS was performed in syncytia one hour later. In order to maintain the same syncytial position for post-stimulation measurements, 2x cAMP-stim buffer (50 μM forskolin, 200 μM IBMX in Fluorobrite DMEM) was added directly to the imaging dish in equal volume to the residual media and a second FCS recording was initiated 5 minutes later. PKA data was fit to a two-component diffusion model:

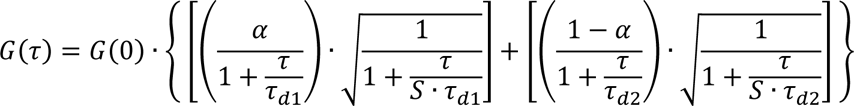

where τ_di_ are the diffusion times for the respective fractions (i = 1, 2), α is the fractional contribution of the first component and all other parameters are as defined above.

### Single particle tracking of membrane proteins in giant syncytia

Membrane proteins are typically free to diffuse in the plasma membrane unless tethered to larger intracellular structures. To examine protein mobility in giant syncytia, U2OS cells expressing mNG-ADRβ2 were co-plated with non-expressing VSVG cells and immersed in fusion buffer for 5 minutes before returning them to Fluorobrite DMEM and incubating at 37°C for 75 minutes. Cells were imaged at 30 fps on a lab-built TIRF microscope at 37°C with an EMCCD (iXon 887, Andor).

Single molecule trajectories were analyzed using the ImarisTrack module in Imaris Bitplane. The raw data was temporally averaged by five frames to improve signal-to-noise, yielding a substack interval of 150 ms, and SPT parameters (PSF quality and size) were chosen based on the dataset. Spots were detected in each substack frame and trajectories were assembled with an autoregressive motion model. 4580 trajectories lasting longer than five substack frames were generated and included for further analysis. MSD plots were fit to extract single particle diffusion coefficients (MSD = 4Dt), which were distributed exponentially. Trajectories were sorted by their diffusion coefficient, heuristically classified into slow, intermediate and fast motions (bottom, middle and top thirds) and average MSD plots were calculated for each of these fractions.

### mNeonGreen folding efficiency and true oligomer calculations

Our FCS data of mNeonGreen (mNG) monomers and covalent dimers permitted an estimation of the fluorescent protein maturation efficiency, denoted as f. In the case of monomers, misfolded proteins are not detected and thus, the detected brightness-per-particle is equal to the brightness of the monomeric species. For simplicity, we normalize this to 1.

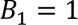

In the case of dimers, we observe a mixture of monomers and dimers. The overall brightness-per-particle (count rate divided by average number of particles) is given by:

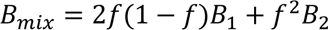

where the prefix to each term denotes the probability of an mNG dimer having one or two fully mature fluorescent domains. Assuming B_2_ = 2B_1_ = 2, this simplifies to:

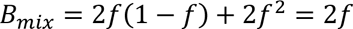

Using our normalized value for B_mix_ (1.7, the observed count rate for dimers divided by count rate for monomers), we estimate that mNeonGreen folds to 85% efficiency.

We can use this fraction to estimate the true dimer propensity of ADRβ2, which was largely present in monomeric and dimeric forms. If we denote the true monomer and dimer fractions as χ_1_ and χ_2_, then we obtain the relation

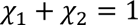

assuming no higher-order oligomers exist.

The “emitter fractions” χ_i_^’^ are given by considering that each ADRβ2 oligomer can have properly folded or misfolded mNG domains. The fractions are given by:

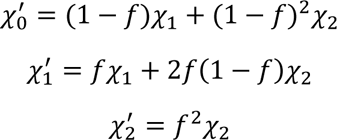

It is important to note that these emitter fractions are not the directly observed quantity (largely because χ_0_ cannot be detected). Instead, we detect

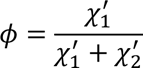

For ADRβ2, φ = 0.75. From the above system of linear equations, we can uniquely determine the values of the “true” monomer and dimer fractions, χ_1_ and χ_2_ (0.64 and 0.36 for ADRβ2).

### Generalized Dark Fraction Correction

Assuming a constant fluorescent protein dark fraction independent of oligomerization state, the observed fractions of each state can be corrected for the presence of dark protein components as an estimate of the actual fraction.

p = fraction of fluorescent proteins (0 to 1.0)

1 - p = dark fraction

The fractional losses of each n-mer state are given by binomial coefficients. These form a series of linear equations that are solved to correct the observed fractions. The matrix form, written here for a maximum oligomer state of 6 (k):

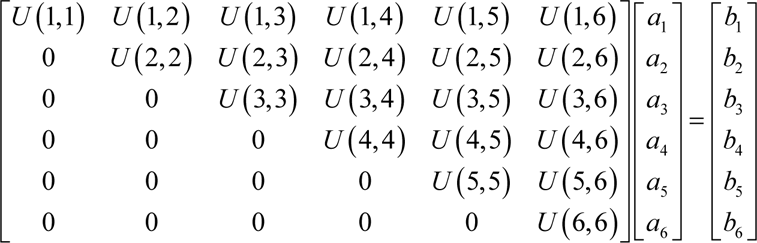

Where the upper triangular matrix (U) elements are:

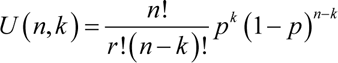 *n* ≡ number of subunits k ≡ subunit number (1 to max assumed)

The vector *b* is the observed number of steps (oligomer subunits) and the vector *a* is the corrected number of subunits per complex. *Ua = b* can be solved for *a* by back-substitution since all of the diagonal elements are non-zero.

### Analysis software for bleach step quantification

#### ImageC

ImageC.exe is a lab-written image analysis program that has single molecule/centroid localization functions useful for single molecule analysis. The application is a Windows based program written in C/C++ using Microsoft Visual Studio 2017. For a portion of the bleach step analysis used in this work ImageC was used to two modes: (1) user-determined number of steps based on observation of time traces, where the users tally the results within a program spreadsheet, or (2) an automated bleach step-counting algorithm that is described in Figure S7.

##### Spot location method

Single molecule spots are located by analysis of an image created by summation of subset of the first 20-50% of image frames. A histogram of the summed image is calculated and pixels at the maximum pixel value located, analyzed either on a peak vs background levels basis for the NxN box (typically 5×5 pixels), or using a Gaussian mask based merit function described below. Once the bright center pixel is analyzed, the pixels within the NxN box are set to zero. The levels criteria asks whether the peak pixel is greater than the user-set background value (BG) for all the pixels within the NxN box. The Gaussian difference mask method compares the NxN pixels surrounding the peak pixel to see if they conform to a Gaussian profile relative the center of the spot. The extent of conformance a user-selectable value that is the fractional ±amount the pixel can deviate from the expected value, based on what would be expected if the spot were a perfect Gaussian. If a spot has nearest neighbor pixels that do not meet this criteria the spot is not used.

##### Gaussian Difference Mask (GDM) method

Assume a Gaussian PSF with a 1/e radius of σ centered at pixel i = j = 0. The camera pixel size is p, and p and σ are in the same units. The pixels at and near the center of the PSF would have values of:

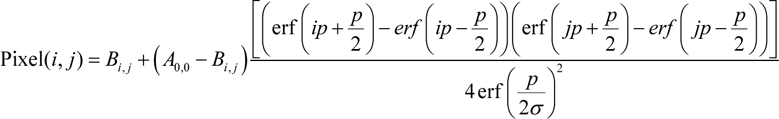

The above can be used to pre-calculate a mask for a given σ and pixel size, which can then be applied as a merit function to judge whether a spot on an image is a PSF:

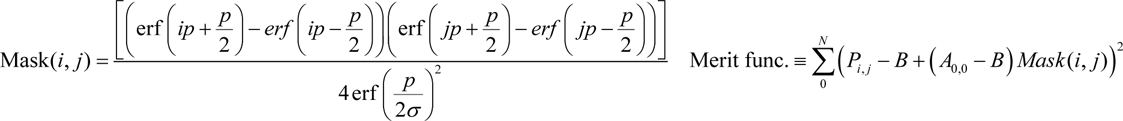

### PIF (ref (36))

In brief, spots are located by scanning a Laplacian of a Gaussian kernel across the surface. Fluorescent puncta above a certain threshold with a good Gaussian fit are selected as valid spots for analysis. The intensity vs time traces of these spots is analyzed using an algorithm which locates intensity drops larger than a predetermined threshold in the trace. A level is determined as the average value of the trace between two drops. The intensity difference between adjacent levels is then compared and they are combined if it is below the threshold. This is repeated in an iterative manner until all remaining levels are separated by an amount greater than the threshold. Each image contained ∼600 spots.

## SUPPLEMENTAL FIGURES

**Figure S1:**
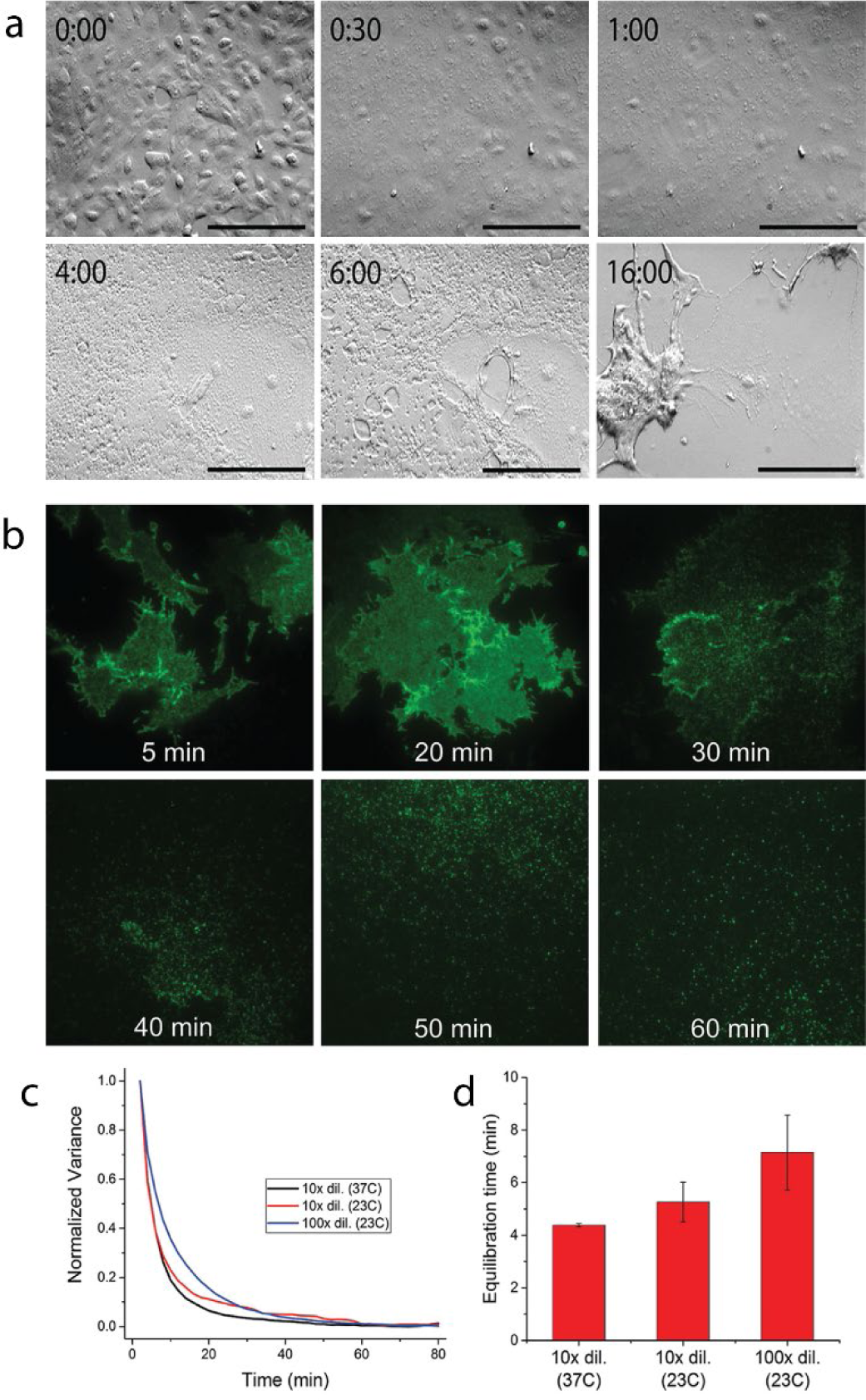
Time course of syncytial formation and protein diffusion in the cytosol and on the membrane. (a) The cell fusion process acquired using an incubator microscope. Initially each cell’s bounding membranes are clearly discernable but this morphology disappears within 30 minutes. The resulting syncytium remains bound to the coverslip for 4-6 hours, at which point it begins to detach from the substrate. By about 16 hours, the syncytium is mostly detached and cells begin to die. Scale bar is 100 μm. (b) mNeonGreen diffusion in the VSVG-induced syncytium measured by the change in signal variance across the image. Equilibrium is reached within 30-60 minutes for all conditions (different co-plating ratios and temperatures). (c) Exponential decay times for image variance in syncytia. Equilibration takes longer for higher co-plating ratios and lower temperatures, as expected for diffusion of macromolecules. (d) Time course of the membrane protein (mNG-ADRβ2) diffusion after cell fusion. Early time points (5 and 20 minutes) show proteins near their original cells. By 30 minutes, single protein complexes are clearly resolved and many fluorescent puncta are visible by 40 minutes.

**Figure S2:**
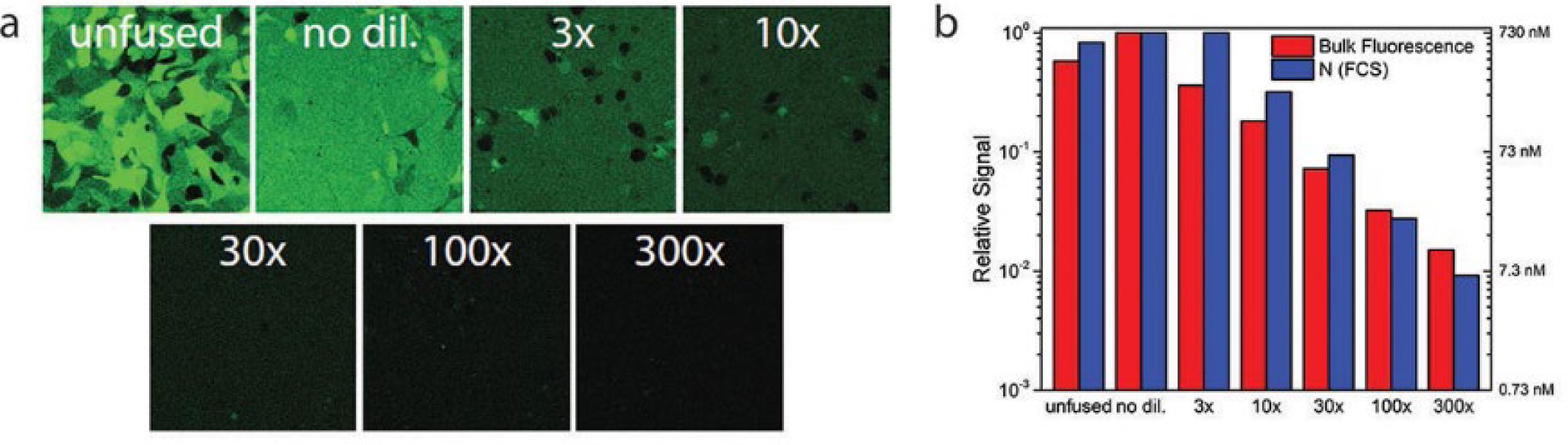
The equilibrium concentration of labeled proteins in syncytia can be set by adjusting the co-plating ratio. (a) A uniform concentration of labeled proteins can be varied over two orders of magnitude with an upper limit constrained only by the number of cells in the culture dish and the diffusion time of labeled protein complexes compared to syncytium lifetime. We find a 1:10 labelled to VSVG cell ratio produces optimum concentrations for single molecule experiments. (b) Quantification of the data shown in (a) using pixel values (red) and fluorescence correlation spectroscopy (blue). FCS permitted calculation of absolute concentrations.

**Figure S3:**
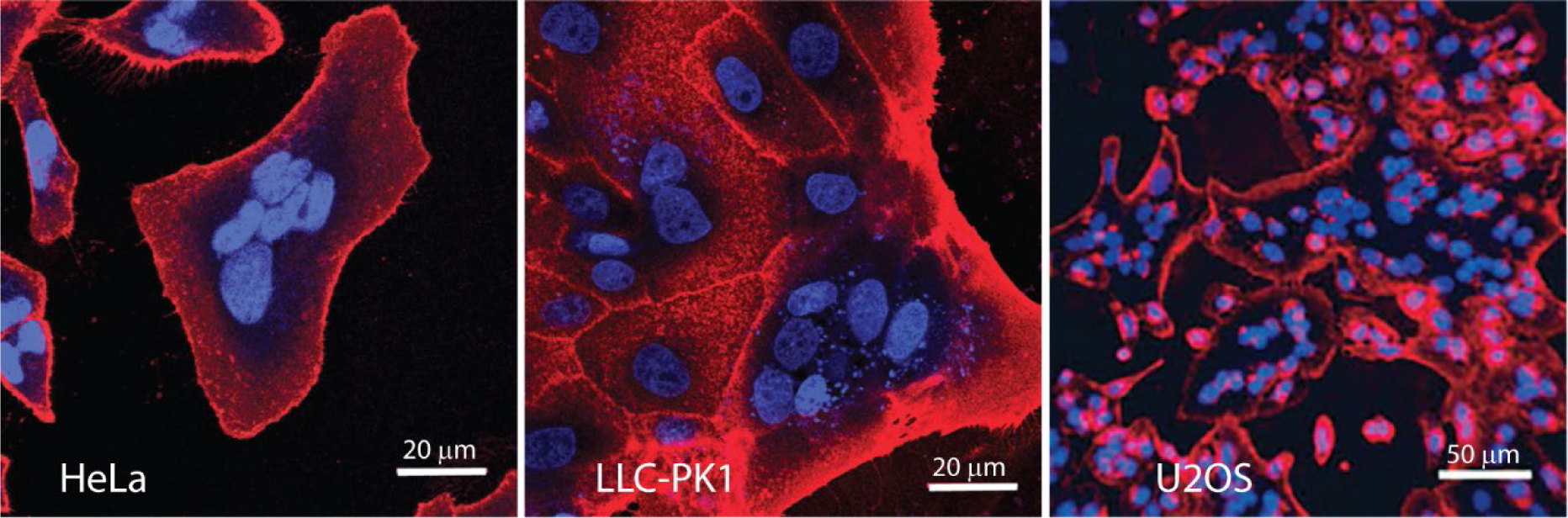
VSVG-mediated fusion in mammalian cell lines. Due to its very broad tropism, VSVG is capable of fusing a large variety of mammalian and non-mammalian cell lines. Here, we demonstrate formation of large syncytia in a human epithelial cancer cell line (HeLa), porcine epithelial cells (LLC-PK1) and Human Bone Osteosarcoma Epithelial Cells (U2OS). Cells were transiently transfected with a constitutive VSVG plasmid (driven by a CMV promoter) and fusion was triggered 24 hours later by a brief incubation in fusion buffer (pH 6.0). After one hour, cells were fixed and membrane and nuclei labeled using Wheat Germ Agglutinin and Hoeschst. Large-scale fusion on similar time scales was also observed in a hamster fibroblast line (BHK-21), human embryonic kidney cells (HEK-293) and Rat Basophilic Leukemia cells (RBL-2H3), a mast cell model (data not shown).

**Figure S4:**
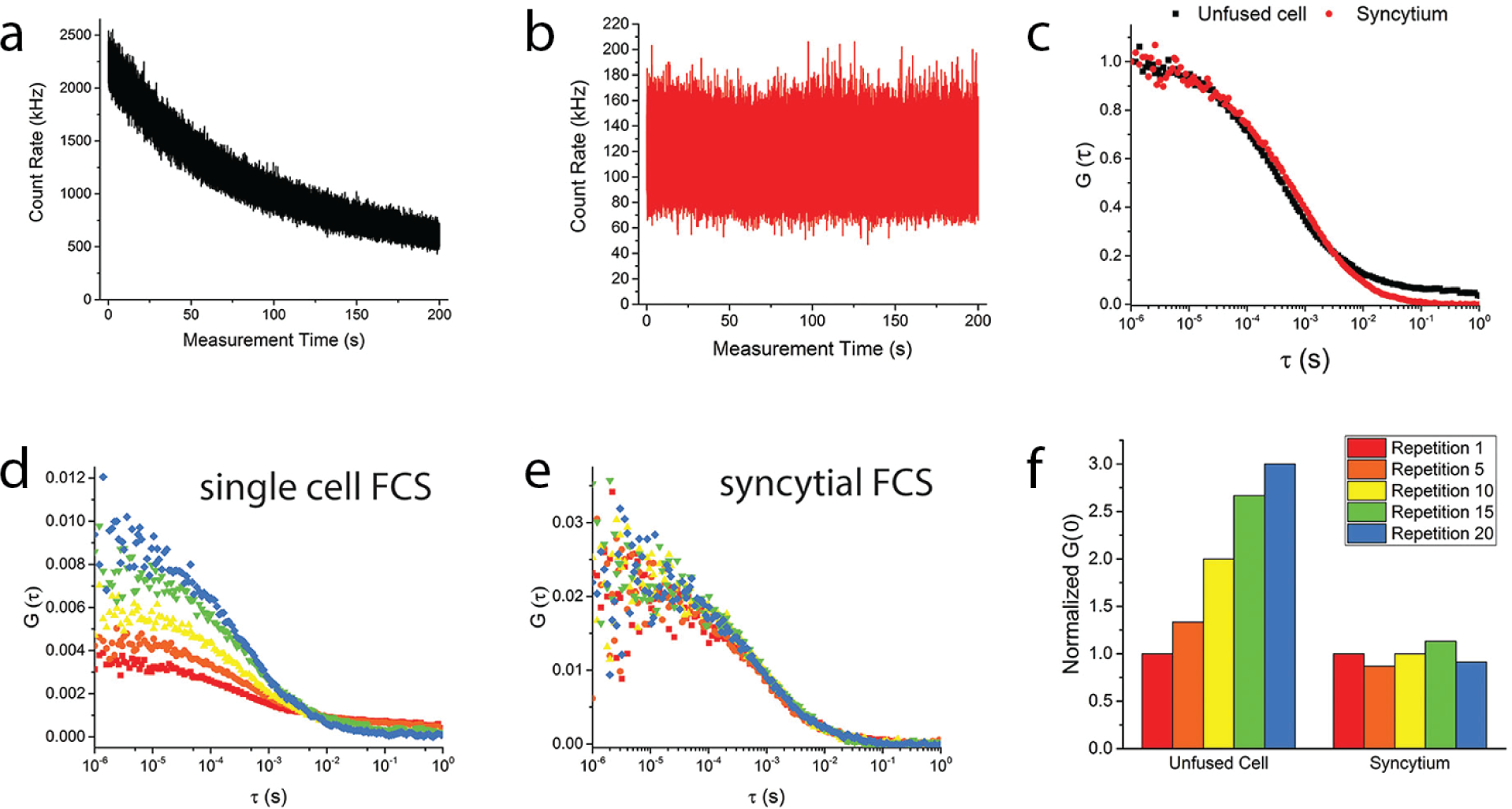
Reduction of live-cell FCS artifacts by the use of SPReAD. (a) Time trace of fluorescence fluctuations in a cell expressing cytoplasmic mNeonGreen. Due to the limited amount of mNeonGreen in a single cell, the effects of photobleaching can be significant. (b) Photobleaching effects are greatly reduced in large syncytia, where the pool of fluorophore is much larger. (c) Correlation curves for the traces in (a) and (b). The correlation curve for mNeonGreen in syncytia fits well to a 1-component diffusion model, while the single-cell correlation shows a poor fit and does not reach its asymptote as expected (due to photobleaching artifacts). (d) Photobleaching in live cells causes the correlation curve to change over time; G(0) continually rises as fluorescent molecules are depleted. (e) In contrast, syncytial FCS shows no time-dependent changes due to the large pool of freely diffusing labeled proteins. (f) Comparison of G(0) variation over time in live cells and syncytia. G(0) systematically rises in single cells while remaining constant (besides noise) in large syncytia.

**Figure S5:**
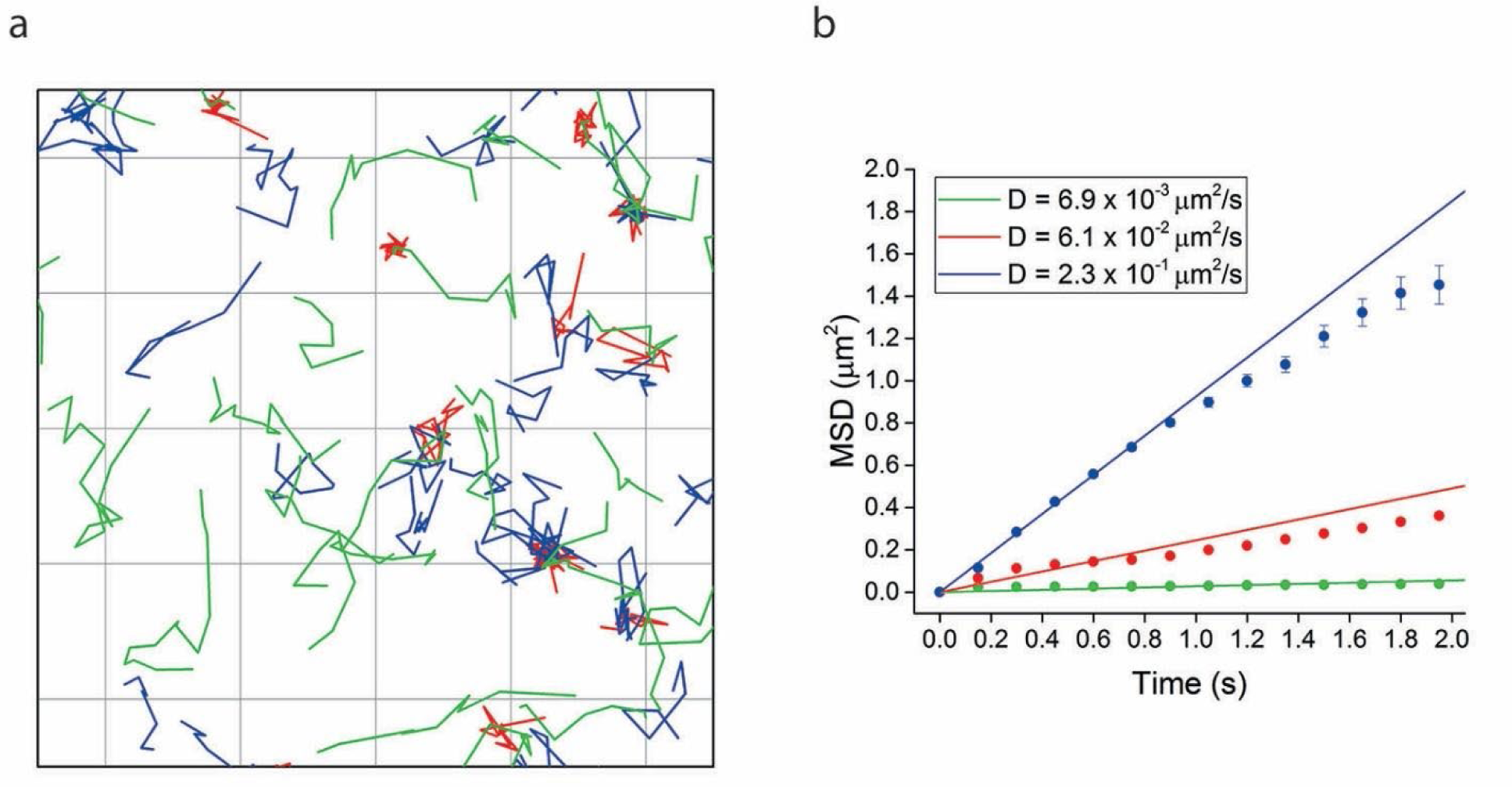
Single-particle tracking of ADRβ2 in large syncytia. (a) Single particle tracking of mNG-ADRβ2 in a 5 μm square of syncytial membrane. Slow, intermediate and fast trajectories show distinct kinetics. Membrane protein diffusion after cell fusion matches previous SPT measurements conducted in live cells, signifying that the syncytial environment preserves the biophysical properties of the plasma membrane. (b) Average MSD plots for slow, intermediate and fast trajectories. Extracted diffusion coefficients agree with the range of parameters measured in live cells.

**Figure S6:**
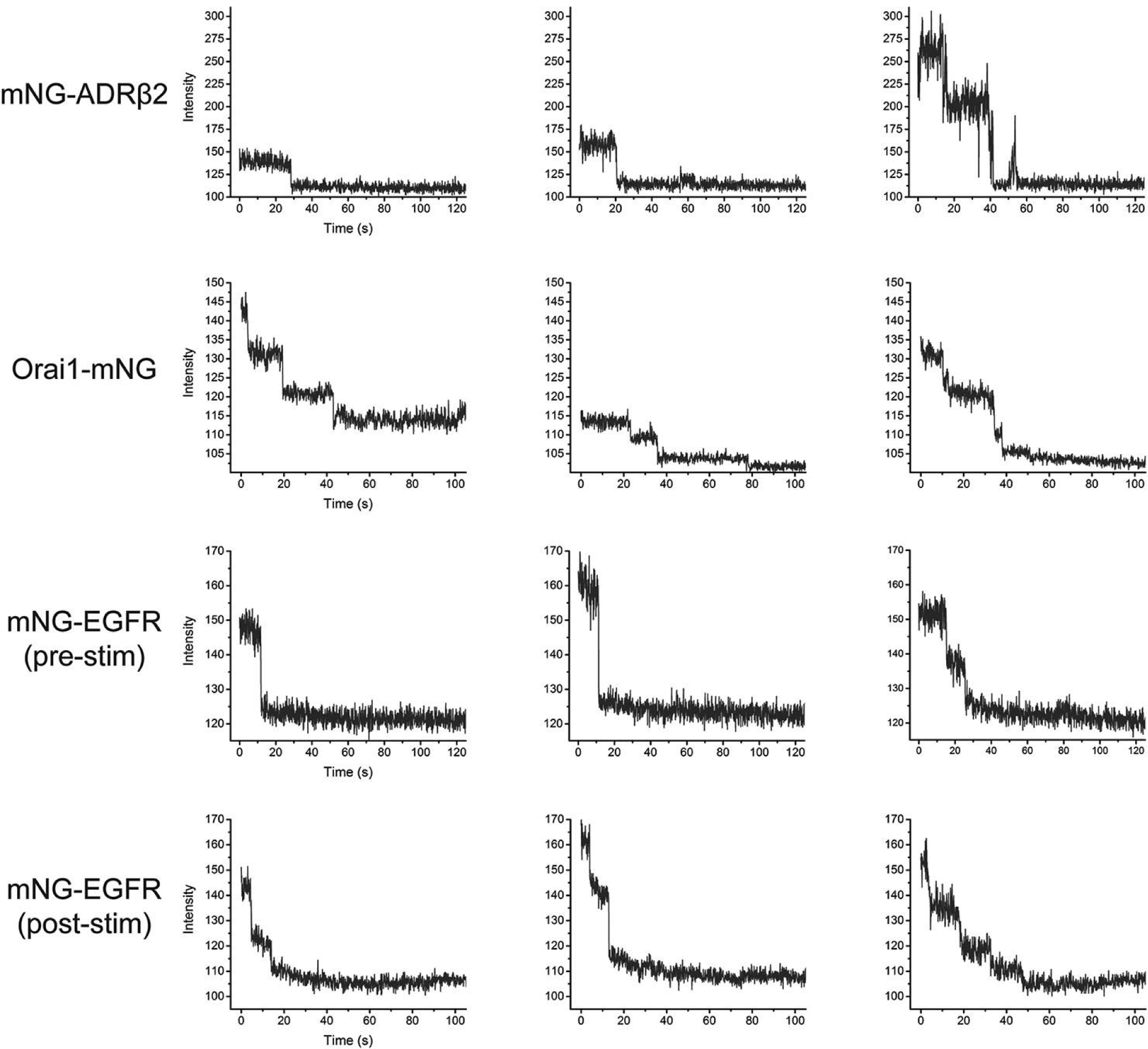
Typical Stepwise photobleaching traces. Sample bleach traces from single protein complexes. Number of photobleach steps are listed below (from left to right): mNG-ADRB2: 1, 1, 2; Orai1-mNG: 3, 3, 4; mNG-EGFR (pre-stim): 1, 1, 2; mNG-EGFR (post-stim): 2, 2, 4

**Figure S7:**
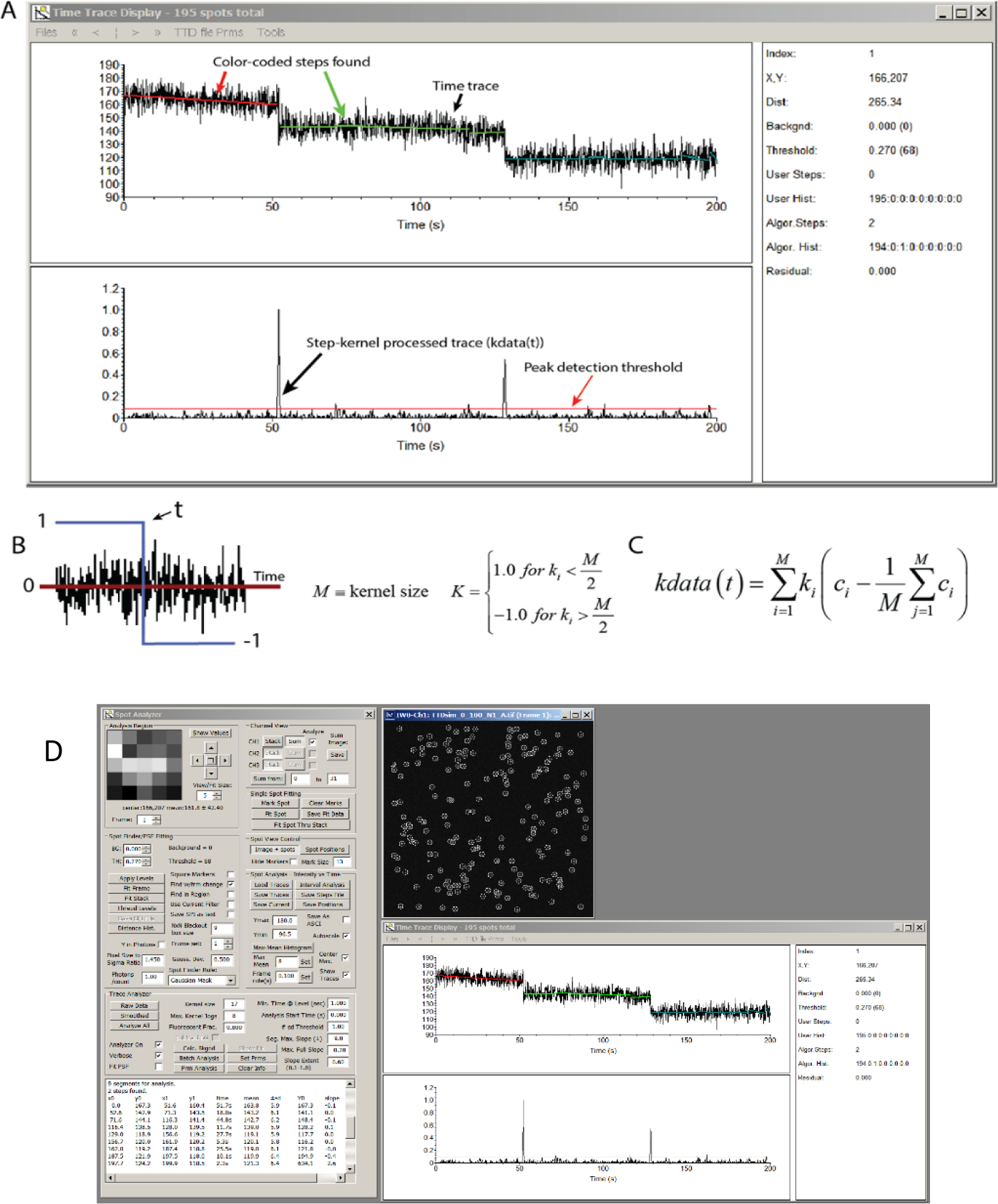
ImageC Analysis software used for bleach step quantification. Bleach Step analysis algorithm initial processing kernel. The algorithm begins by convoluting the time trace with the kernel defined in B. kdata(t) is effectively the derivative of the time trace. kdata(t) is squared to make it positive, normalized and displayed in the lower plot window of the Time Trace Display window. The peaks in kdata(t) locate potential bleach steps in time trace as shown above in A. The peak detection threshold level determines how many steps (referred to as “Kernel Jogs” in the software) to use in the processing algorithm (Max. Kernel Jogs parameter in Figure 7). Once a peak threshold is found that found produces the (user-set) number of allowed jogs, the time trace is broken up into #jogs + 1 segments defined by the threshold crossing points. The segments are fit to a linear function to obtain their slope. A characteristic deviation (noise level) is calculated for each segment and if the segments are of acceptable length (in time) and slope, they are further analyzed to identify whether they overlap in pixel value. If so, they are considered to be in the same level group. The total number of levels are tallied and they number – 1 (to account for the final fully bleach level) is reported by ImageC to be the number of photobleach steps in the trace.

**Figure S8:**
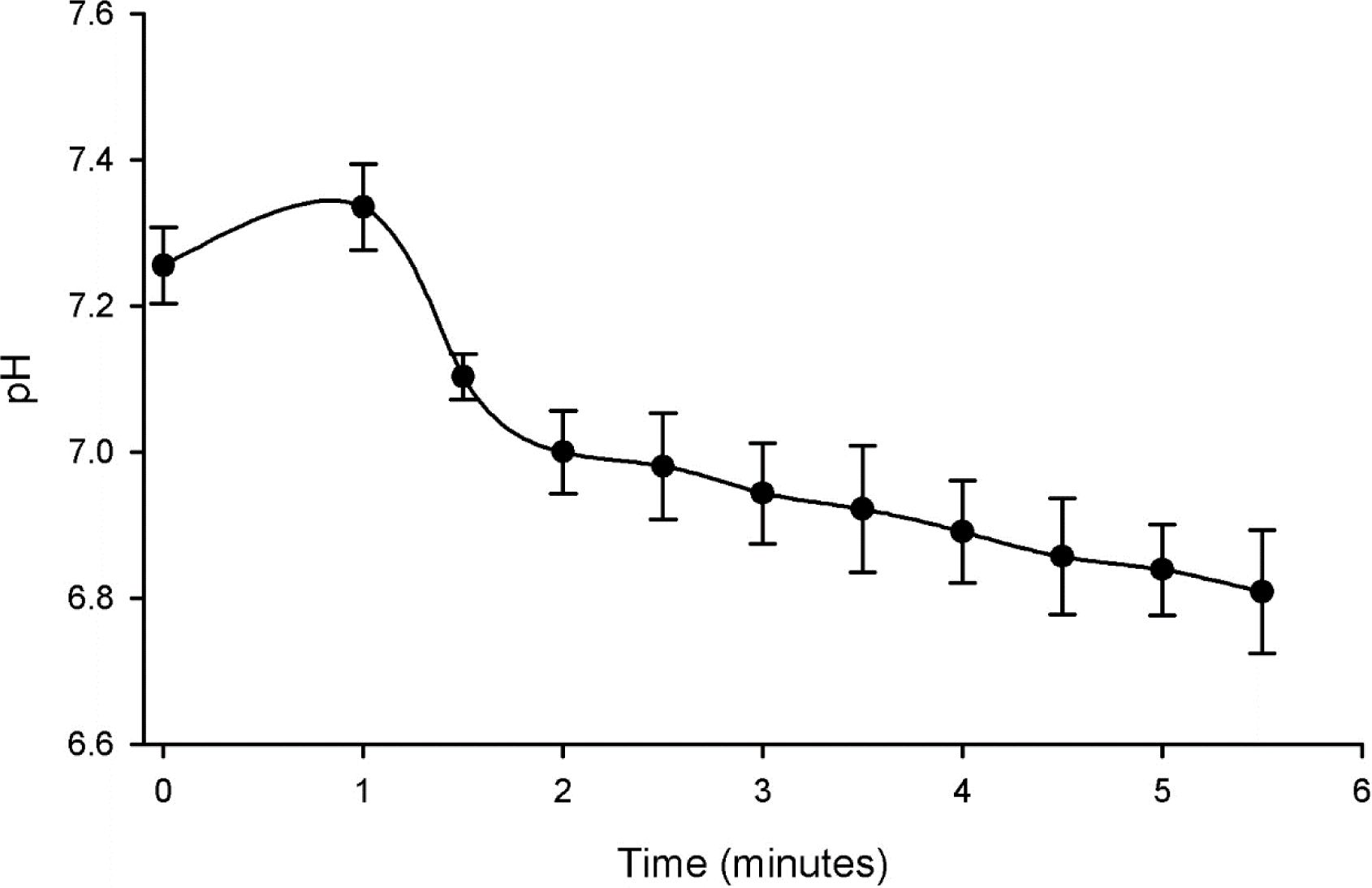
Effect of the short pH drop on intracellular pH using SNARF. After a 24 hr induction period in doxycycline, Tet-VSVG U2OS cells fuse immediately after a brief pH drop from 7.4 to a pH < 6.0. We measured the effect on the intracellular pH during incubation at pH 5.5 and found that during the short time needed for activating VSVG (0-2 minutes) the pH remains above pH 7.0. Even during longer incubations (up to 5 minutes), the pH was never found to be lower than ∼6.8. Data are mean ±SEM n = 2.

**Table S1:**
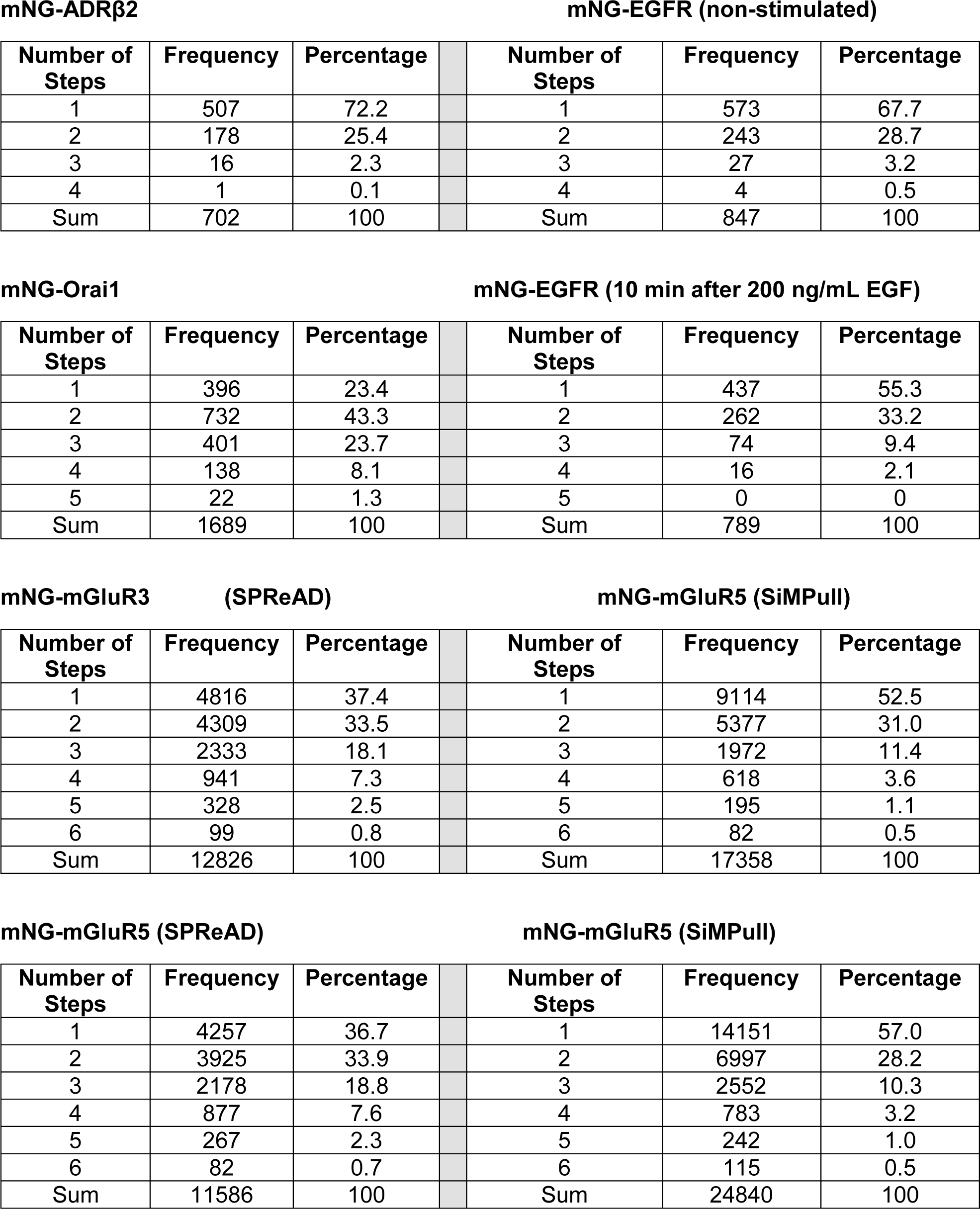
Photobleach step histograms Frequency of bleach steps for each of the membrane protein oligomers studied.

## Supplementary Movies

**Video 1: mNG-ADRβ2 mobility after cell fusion (ADRBeta2_MobilityInFusedCells.avi)** Diffusion kinetics of mNeonGreen-tagged ADRβ2 oligomers in syncytial membranes are similar to membrane proteins in unfused cells, suggesting that cell fusion does not significantly perturb the biophysical character of the plasma membrane.

**Video 2: Long time course imaging of syncytial formation (FusionTimeCourse_Brightfield.avi)**

Oblique-illumination brightfield imaging after pH drop. Cells fuse at beginning of movie and bounding membranes are seen to disappear within 30 minutes. A single syncytium remains bound to the imaging dish for 5-6 hours, after which it begins to detach from the glass coverslip. Detachment progresses until ∼20 hours, at which point massive cell death is observed. Fluorescence correlation spectroscopy or formaldehyde fixation (for TIRF experiments) shown is this work were performed during the first 1-2 hours to minimize perturbation of cellular protein complexes.

**Video 3: Confocal imaging of cytoplasmic protein diffusion during cell fusion (mNeonGreen_InFusingCells.avi)**

Dynamics of mNeonGreen diffusion out of parent cells and into the larger syncytium. Cytoplasmic fluorescent proteins are mobile immediately following pH drop, with equilibrium concentrations being reached within 30-40 minutes (for cytoplasmic mNeonGreen). Equilibration time depends on the molecular weight of the diffusive species, interactions with other cellular components, the ratio of expressing:non-expressing cells and temperature which was 10 to 1 in this case. Field of view is 40 x 40 μm.

